# Differentiation of human induced pluripotent stem cells into functional airway epithelium

**DOI:** 10.1101/2020.11.29.400358

**Authors:** Engi Ahmed, Mathieu Fieldes, Chloé Bourguignon, Joffrey Mianné, Aurélie Petit, Charlotte Vernisse, Myriam Jory, Chantal Cazevieille, Hassan Boukhaddaoui, James P. Garnett, Gladys Massiera, Isabelle Vachier, Said Assou, Arnaud Bourdin, John De Vos

## Abstract

**Rationale:** Highly reproducible *in vitro* generation of human bronchial epithelium from pluripotent stem cells is an unmet key goal for drug screening to treat lung diseases. The possibility of using induced pluripotent stem cells (hiPSC) to model normal and diseased tissue *in vitro* from a simple blood sample will reshape drug discovery for chronic lung, monogenic and infectious diseases.

**Methods:** We devised a simple and reliable method that drives a blood sample reprogrammed into hiPSC subsequently differentiated within 45 days into air-liquid interface bronchial epithelium (iALI), through key developmental stages, definitive-endoderm (DE) and Ventralized-Anterior-Foregut-Endoderm (vAFE) cells.

**Results:** Reprogramming blood cells from one healthy and 3 COPD patients, and from skin-derived fibroblasts obtained in one PCD patient, succeeded in 100% of samples using Sendai viruses. Mean cell purity at DE and vAFE stages was greater than 80%, assessed by expression of CXCR4 and NKX2.1, avoiding the need of cell sorting. When transferred to ALI conditions, vAFE cells reliably differentiated within 4 weeks into bronchial epithelium with large zones covered by beating ciliated, basal, goblets, club cells and neuroendocrine cells as found *in vivo*. Benchmarking all culture conditions including hiPSCs adaptation to single-cell passaging, cell density and differentiation induction timing allowed for consistently producing iALI bronchial epithelium from the five hiPSC lines.

**Conclusions:** Reliable reprogramming and differentiation of blood-derived hiPSCs into mature and functional iALI bronchial epithelium is ready for wider use and this will allow better understanding lung disease pathogenesis and accelerating the development of novel gene therapies and drug discovery.

## Introduction

Chronic obstructive pulmonary disease (COPD), is one of the leading causes of death worldwide [1]. Induced pluripotent stem cells (iPSCs) represent an attractive opportunity compared with the existing solutions to model chronic airway diseases because they can yield a virtually unlimited amount of any differentiated cell type [2]. The recent description of protocols to differentiate human pluripotent stem cells (PSCs) into bronchial epithelium has been encouraging [3–11]. Overall, these protocols rely on the knowledge gathered on normal lung development in mammals [12]. Briefly, lung embryogenesis starts with the formation of the definitive endoderm (DE). During the 4^th^ week of human embryonic development, the primitive gut appears and can be divided into foregut, midgut, and hindgut. Early pulmonary development starts from the ventral area of the anterior foregut endoderm (vAFE). From this zone, which is characterized by the expression of the transcription factor NKX2.1, the respiratory diverticulum will emerge and form the trachea, and then bronchi, bronchioles, and alveoli. These steps can be recapitulated *in vitro* by differentiating PSCs first into DE and then by driving DE cells towards vAFE differentiation [13]. Finally, vAFE cells are specifically differentiated into lung progenitors and then bronchial cells. However, the protocols for PSC differentiation into bronchial epithelium present several limitations, and multiplicity of protocols were rarely described in detail. Most of them are effective on a very limited number of cell lines, most often healthy control cells and require an enrichment step based on a specific NKX2.1+ cell selection at the vAFE stage using flow cytometry and cell surface markers (e.g. carboxypeptidase M (CPM)+ cells [9] or CD47^hi^CD26^lo^ cells [14]), or a final differentiation step in 3D culture conditions. Others require important technical skills and are difficult to replicate [15].

Here, we developed an approach to differentiate human iPSCs (hiPSCs) into proximal airway epithelium, using a straightforward protocol without any cell purification step. Careful in-home reprogramming and then culture adaptation to single-cell passaging together with a precise timing and reagent benchmarking for each differentiation step led to the successful generation of fully differentiated and functional bronchial epithelium in air-liquid interface (ALI) culture conditions from hiPSCs (iALI bronchial epithelium). We successfully used this protocol to differentiate five hiPSC lines, among which three were derived from patients with severe COPD. This study highlights the criticality of evaluating expansion and differentiation conditions for achieving optimal phenotypic and functional endpoints such as ciliary beat frequency (CBF), mucus flow velocity, presence of differentiated cells, transepithelial electrical resistance (TEER). This simple protocol to produce hiPSC-derived bronchial epithelium in ALI culture conditions (iALI bronchial epithelium) will facilitate modelling airway diseases developing novel gene or cell therapies, and drug discovery.

## Results

### Reprogramming from a blood sample or skin-derived fibroblasts

Skin-derived fibroblasts from the patient with primary cilia dyskinesia (PCD) [16] or peripheral blood mononuclear cells from the healthy control and the three patients with severe COPD and were reprogrammed using Sendai virus to generate the PCD02.30, HY03, iCOPD2, iCOPD8, and iCOPD9 hiPSC lines, respectively (figure 1). From venepuncture, PBMC were Ficoll-isolated and cultured using STEM SPAN SFEMII ® kit enriched with cytokines (IL3, SCF, EPO) promoting Erythroid Progenitor (EP) expansion. CD45, CD34, CD71 and CD36 monitoring were required to optimize yield of EP expansion before Sendai virus transduction. C-myc, KLF4, SOX2 and OCT4-containing Sendai viruses were concomitantly added once to the EP culture for three days. After transfer into Geltrex, hiPSC clones were observed 30 days after blood sampling. Pluripotency was confirmed by demonstrating phosphatase alkaline activity, cell surface SSEA3/4 and TRA1-60 expression, and OCT4, NANOG, SOX2 mRNA expression. HiPSC genetic integrity was assessed by ddPCR (iCS digital) (supplemental figure 2A and B) [18]. One of the COPD-reprogrammed iPSC clones (iCOPD2) was found to harbour one genomic abnormality, copy number gain in 20q11.21, yet differentiation could still be achieved with this clone.

**Figure 1:**
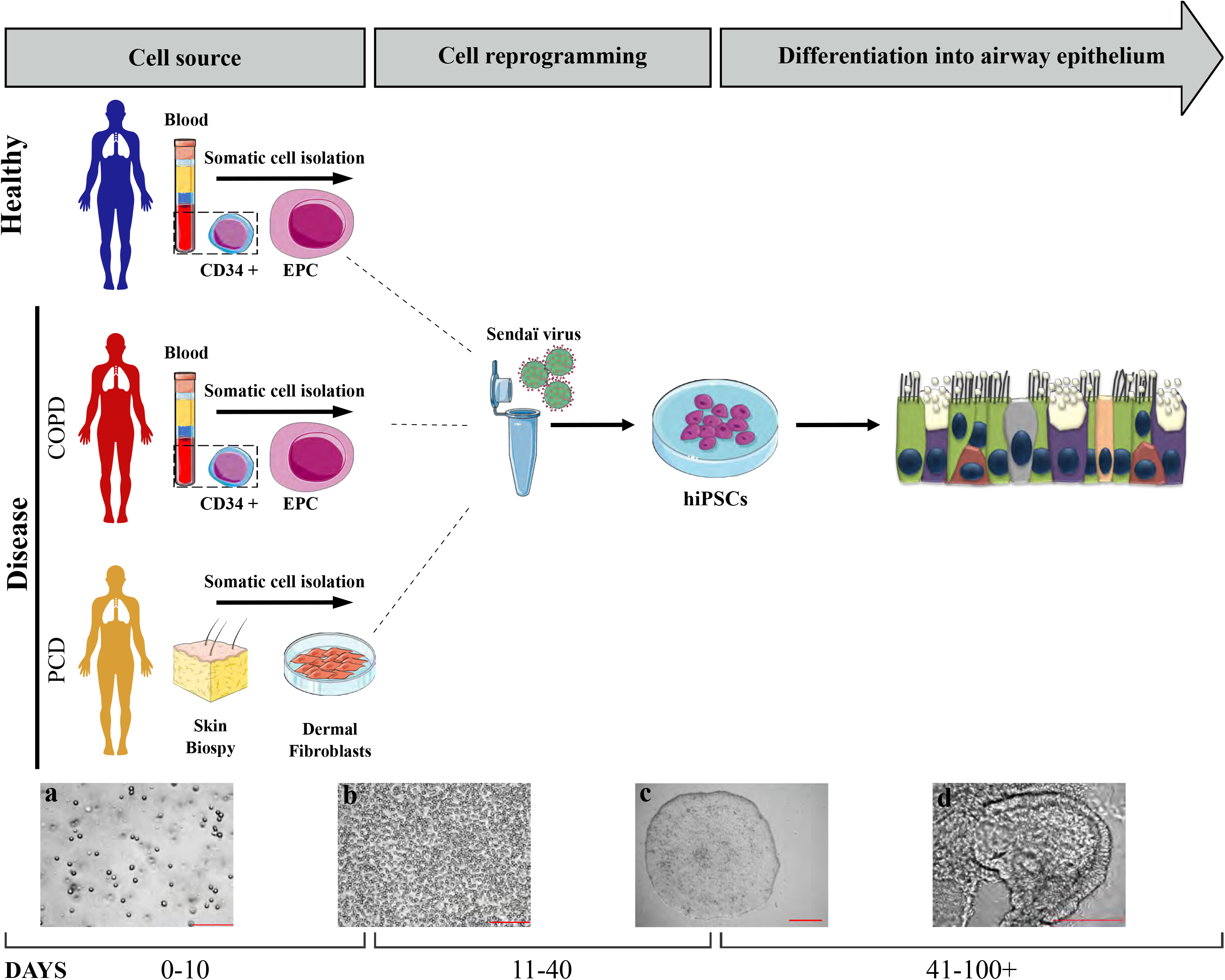
Workflow of the study protocol: from iPSC generation to iPSC-derived airway epithelium. Left panel: recovery of cell source Day 0 to Day 10. Peripheral blood mononuclear cells (PBMC) (a) were isolated from whole blood sample from healthy and COPD patients. CD34+ subpopulation (b) was amplified into erythroid progenitor cell (EPC). Fibroblasts were isolated from a skin biopsy of a PCD patient and amplified *in vitro*. Middle panel: Cell reprogramming step Day 11 to 40. EPC or fibroblast were transduced using Sendai virus constructs containing Oct3/4, Sox2, Klf4 and c-Myc. Induced pluripotent cell (IPS) colony were visible at Day 40 (c). Right panel: iPSC differentiation into airway epithelium, from day 41 to Day 100+ (d). COPD: chronic obstructive disease; PCD: primary ciliary dyskinesia, EPC: erythroid progenitor cell.

### Adaptation of hiPSCs to single-cell culture is mandatory for a successful differentiation process and allows high rate of definitive endoderm induction

The differentiation protocol, is schematized in figure 2A. To develop a robust differentiation protocol, we benchmarked the timing, the cell density and the method of passaging, factors that were crucial for achieving reliable rates of DE purity and quality. hiPSC lines were passaged as single cells because hiPSC clumps were partly resistant to DE induction, as evidenced by OCT4 expression persistence. Optimal cell adaptation was obtained by gentle colony dissociation into small clumps for five passages, and then into single cells for at least 5-10 passages, using Versene (EDTA) in the presence of Y-27632 (figure 2B). Adaptation to single-cell passaging was deemed mandatory to prevent massive cell death after cell plating for APS induction (Figure 2C). Then, the differentiation process was started by adding activin A, and CHIR99021 (a GSK3 inhibitor that acts as a WNT pathway agonist) in the presence of the ROCK inhibitor Y-27632 for one day (day 1; Anterior Primitive Streak figure 2A and supplemental table 3) followed by activin A, LDN-193189 and Y-27632 for 1-2 days, leading to DE induction (day 2-3, figure 3A). To optimize the protocol, various intervals between hiPSC plating and anterior primitive streak (APS) induction, as well as different cell densities (from 70 to 130K cells/cm^2^) were tested (figure 2D-E). Indeed, plating cells at too low density led to important cell death, whereas too high density led to persistent and sustained OCT4 expression (figure 2F). This optimized protocol robustly yielded in average more than 80% of CXCR4+ DE cells within 2-3 days, and was validated using the five hiPSC lines described above and compared with one human embryonic stem cell (ESC) and two other hiPSC lines (n=170 independent experiments, using eight PSC lines) (figure 3A-C and supplemental figure 3). Moreover, DE cells expressed characteristic endoderm transcription factors FOXA2 and SOX17 (figure 3D-E). Pluripotent markers NANOG, SOX2 and OCT4 progressively switched off with the stage progression (figure 4E).

**Figure 2.**
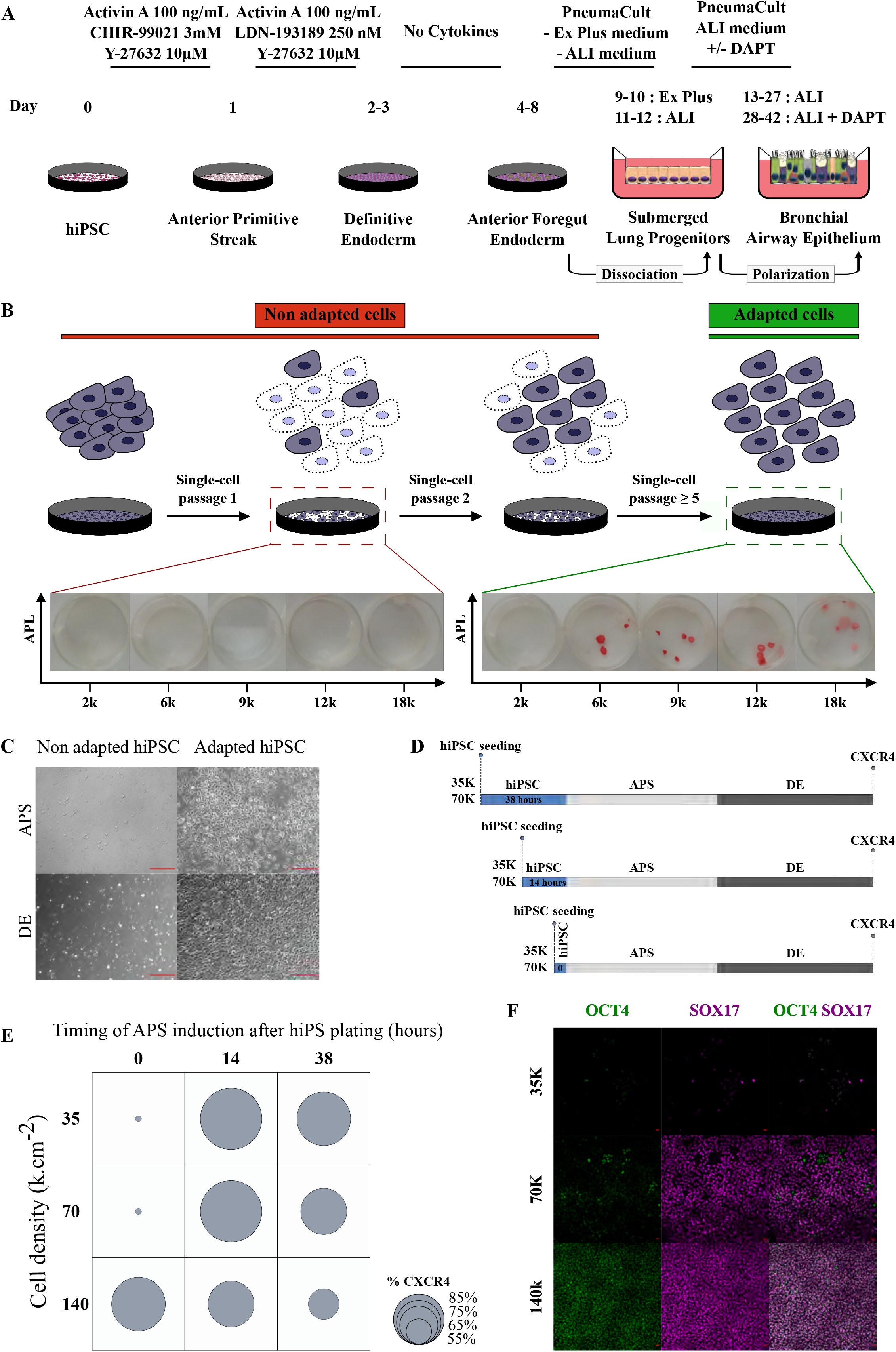
Adaptation to single-cell culture is required before starting differentiation. (A) Schematic representation of the differentiation protocol. (B) Left: Confluent hiPSC colony culture passaged in mechanical clumps. Middle: Non-adapted hiPSC cells (<5 single-cell passages) undergo massive cell death. Right: Adapted cells after serial single-cell passages. APL: Alkaline Phosphatase staining on hiPSC that were plated at low density and grown for one week. Hy03 cell line. (C) Left panels: Non-adapted hiPSCs show massive cell apoptosis at the APS/DE stage. Right panels: Confluent cell layer at the APS/DE step when using adapted cells, Hy03 cell line. (D) Design of the experiments to optimize the interval between hiPSC plating and APS induction (two plating densities: 35 000 and 70 000 cells per cm-2). (E) Results of the optimization experiments based on CXCR4 expression (DE marker). (F) Too low (35K.cm-2) and too high (140K.cm-2) cell plating density lead to massive cell death or incomplete OCT4 inhibition, respectively. Optimal cell density (here 70K.cm^2^) induces strong OCT4 inhibition and high SOX17 expression (iCODP8 cell line).

**Figure 3.**
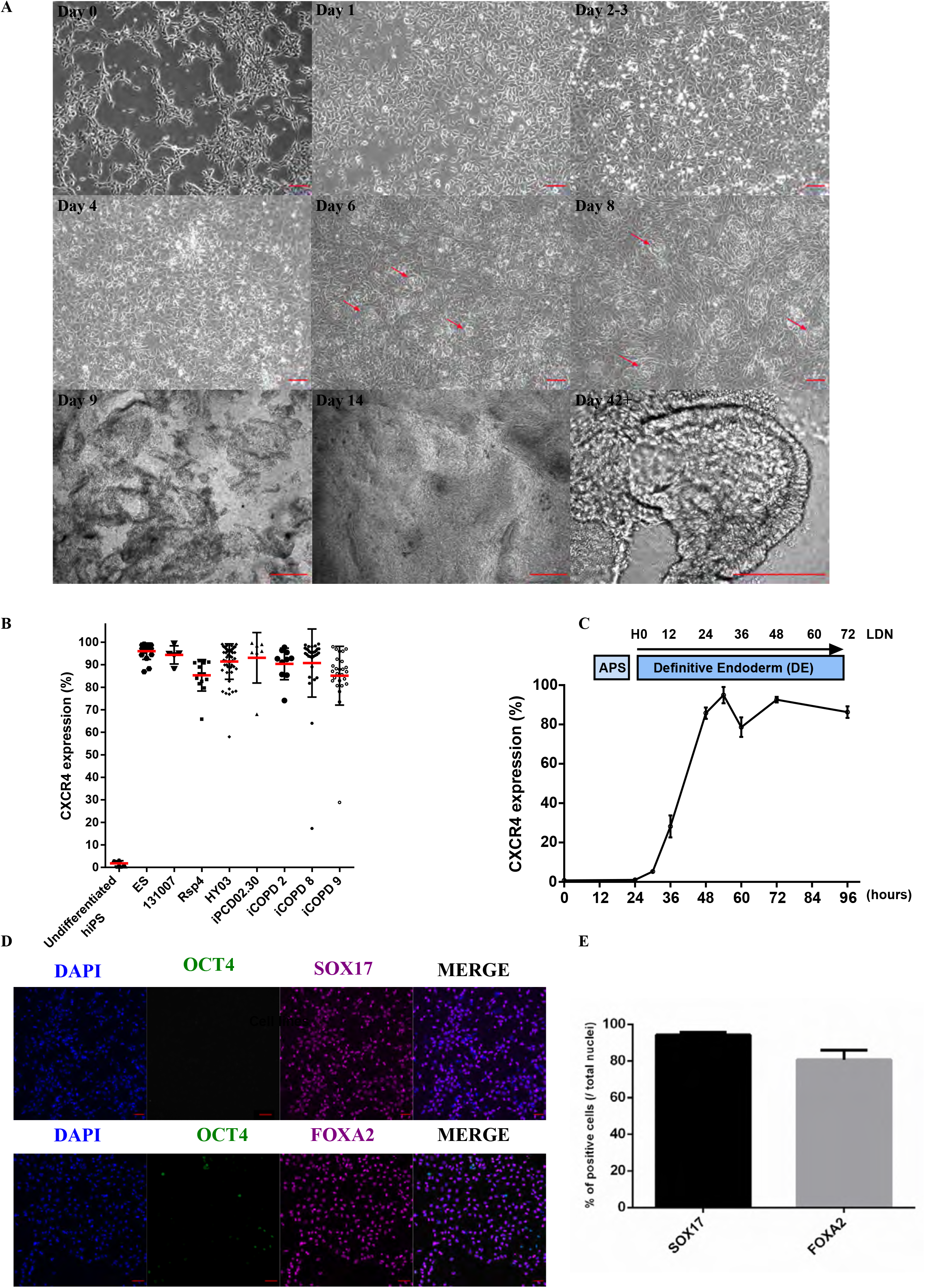
Differentiation of induced pluripotent stem cells into bronchial airway epithelium. (A) Morphological changes during the different differentiation steps. Day 0: hiPSC cells plated as single cells. Day 1: Anterior primitive streak. Day 2-3: Definitive endoderm. Day 4, 6 and8: Anterior foregut endoderm; red arrows: bud-like structures. Day 9: Lung progenitors after mechanical clump passage and plating on Transwell inserts. Day 14 (polarization day): Epithelial layer. Day 42+: Multi-ciliated bronchial epithelial layer. Scale bar 200μm. (B) Quality of DE induction based on CXCR4 expression by flow cytometry analysis in the different PSC lines used (n=8) (C) Time course of DE induction (n=3, HY03 hiPSC line). (D) Immunofluorescence analysis of OCT4, SOX17 and FOXA2 expression in DE cultures derived from PCD cell line. (E) Quantification of SOX17 and FOXA2 positive cells as percentage of all DAPI-positive cells, (n=3, PCD02.30 cell line).

**Figure 4.**
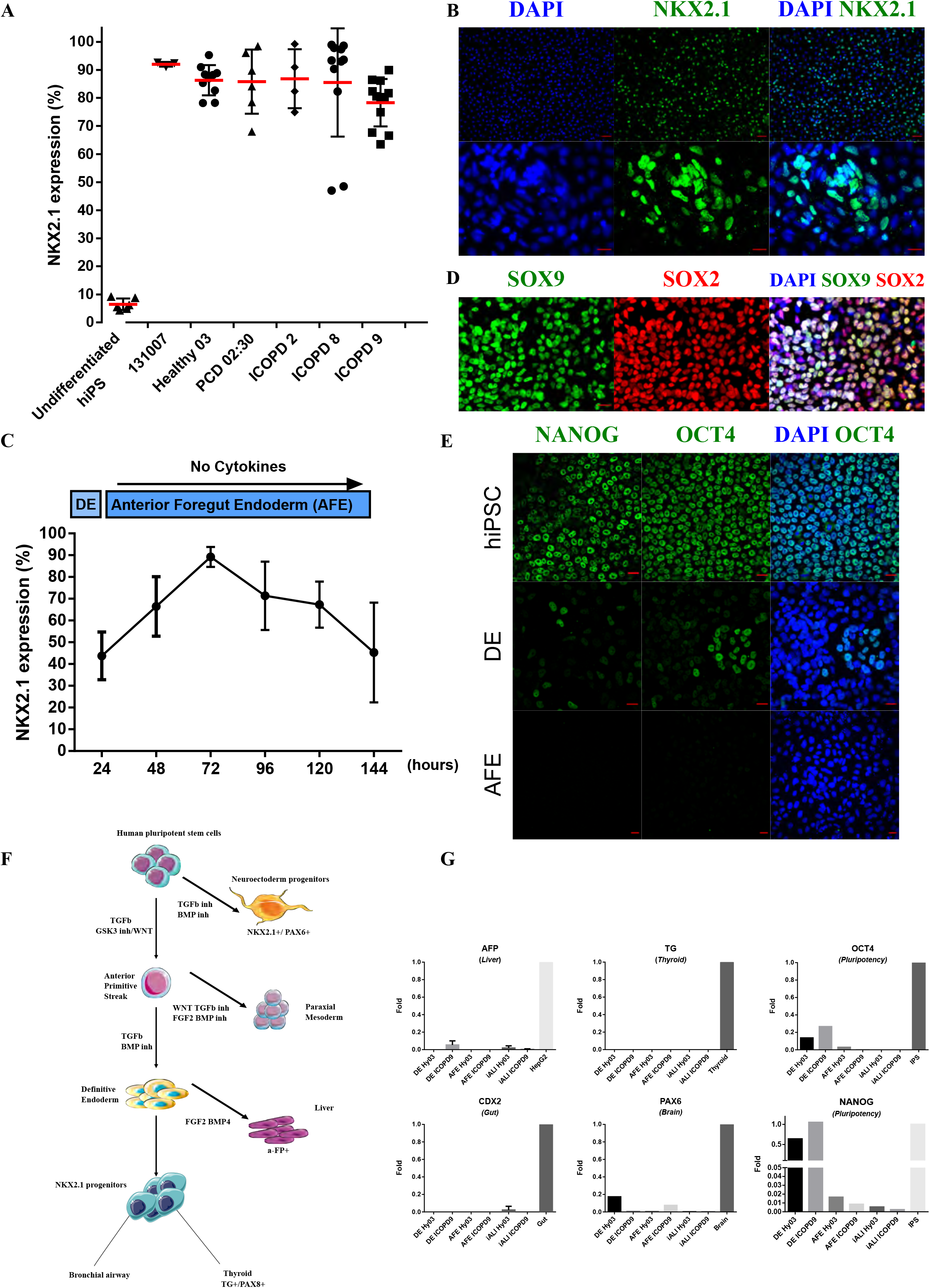
Anterior foregut endoderm characterization. (A) Percentage of NKX2.1-positive cells after vAFE induction in the indicated cell lines. Undifferentiated hiPSCs: negative control. (B) Expression of NKX2.1, a ventral anterior foregut endoderm marker, assessed by immunofluorescence (Hy03 cell line). (C) Kinetics of NKX2.1 expression (n=3, HY03 cell line). (D) Expression of SOX2 and SOX9. Note, the presence of SOX2/SOX9 double-positive cells. (E) Analysis of the pluripotency markers NANOG and OCT4 in hiPSCs (top), definitive endoderm (DE; middle) and ventral anterior foregut endoderm stage (vAFE; bottom) (Hy03 cell line). (F) Model of hiPSC differentiation into the three embryonic layers, emphasizing that NKX2.1 expression in shared by bronchial, neuroectodermal and thyroid progenitors, the two later being a potential source of cell contamination in NKX2.1 positive cells during iPSC differentiation into lung progenitors. (G) Quantitative PCR analysis to assess contamination by thyroid gland (TG), liver (AFP), brain (PAX6) or intestine (CDX2) and confirm progressive extinction of pluripotency marker OCT4 and NANOG. Positive controls: brain mRNA, gut mRNA, thyroid mRNA, HepG2 (human liver cancer cell line) mRNA. IPS used for pluripotency control. Scale bar = 20μm.

### Efficient induction of high purity NKX2.1+ lung progenitors without need for cell sorting

The comparison of various combinations of growth factors for vAFE induction showed that DE cells needed minimal cell signalling, and therefore, were grown in RPMI1640 basal medium with B27 minus vitamin A (figure 2A and Supplemental tables 2, 4 and 5). For efficient vAFE induction, a DE cell population with at least 80% of CXCR4+ cells was required. Time course experiments showed that at 24-36 hours after LDN-193189 addition, there was a narrow window when cells exhibited optimal conditions (i.e. high CXCR4 expression and high viability) for vAFE induction. 3D bud-like structures emerging between days 4-8 appeared to be a good morphological indicator of vAFE differentiation at optical microscopy (figure 3A, red arrows). In these conditions, >80% of cells consistently expressed NKX2.1, as indicated by flow cytometry and confirmed by immunolabelling, in six different PSC lines (n=46 independent experiments) (figure 4 A and B, supplemental figure 3). The optimum percentage of NKX2.1+ cells (>80%) was observed around day 3 after vAFE induction (figure 4C). This result was confirmed by time course immunostaining at v-AFE, with gradual increase of NKX2.1 expression over the time (Supplemental figure 4A). This NKX2.1 expression level was required to induce an efficient differentiation process towards iALI. Interestingly, we were able to detect SOX2, SOX9 expression at protein level by immunostaining. Three populations were observed, SOX2+/SOX9-, SOX2-/SOX9+ and bipotent progenitors SOX2+/SOX9+, as previously reported *in vivo* during human lung development (figure 4D, supplemental figure 4B) [19]. Extinction of pluripotency markers such as OCT4 and NANOG expression were observed at this stage, compared with the DE stage (figure 4E-G). NKX2.1 bronchial progenitor cells exhibited a high proliferation rate, assessed by Ki67 labelling (Supplemental figure 4C). Terminal airway epithelial markers were not detected during v-AFE induction, ascertaining the immature feature of these progenitor cells, consistent with other human iPSC protocol differentiation [14] and *in vivo* mouse lung development [19] [20]. As NKX2.1 is also expressed in other developing tissues (figure 4F), we assessed the purity of NKX2.1 cells by confirming the absence of contamination by RT-qPCR analysis of specific mRNA for thyroid gland (thyroglobulin (TG)) and brain (Paired Box 6 (PAX6)) cell markers, as well as gut (caudal type homeobox 2 (CDX2)) markers (figure 4G). Absence of liver contamination was confirmed at both mRNA and protein level (alpha-fetoprotein (AFP)) (figure 4G, supplemental figure 4D).

### Specification of NKX2.1 lung progenitor cells under 2D ALI culture conditions lead to functional, multi ciliated airway epithelium

iPSC-derived ALI (iALI) bronchial epithelium was obtained from five different iPSC lines (n>3 independent experiments per cell line). v-AFE cells were mechanically dissociated into small clumps and plated at high density on Transwell inserts in PneumaCult-Ex Plus medium (Day 9, figure 2A). At day 2 post-seeding in PneumaCult-Ex Plus medium, cells were progressively switched to PneumaCult-ALI maintenance medium. Four days after seeding on Transwell inserts, medium was removed from the apical side to switch to ALI culture (“polarization”). DAPT, a γ-secretase inhibitor that blocks NOTCH signal transduction, was added to the culture medium present in the basolateral part of the Transwell from day 14 to day 28 post ALI (figure 2A and supplemental table 3).

#### Epithelium with barrier function

iPSC-derived epithelial cells reached confluence after the four days of submerged growth conditions (figure 3A). Features consistent with epithelium could be identified by : i) optical microscopy at late iALI stage (day 42+; figure 5A) ii) immunolabeling of E-cadherin protein (figure 5A) and iii) adherent junctions presence (junctional complexes) assessed by transmission electron microscopy of day 34 post-ALI cultures (Supplemental figure 5A). Barrier integrity of cells during ALI 2D-culture differentiation was assessed by transepithelial electric resistance (TEER). TEER increased significantly during the differentiation process (Supplemental figure 5B), reaching around 300 Ω.cm^2^ and could be maintained for >200 days of culture.

**Figure 5.**
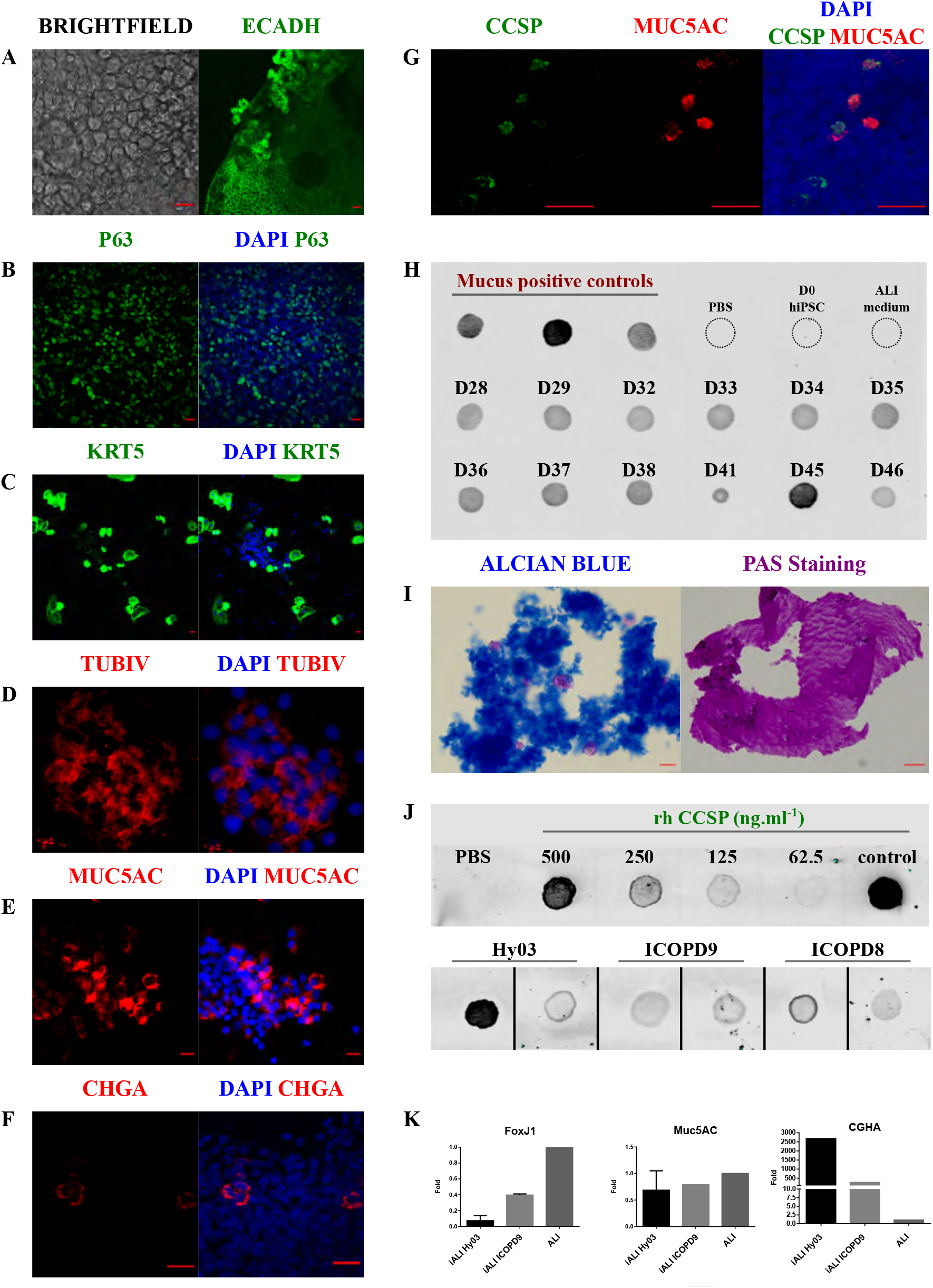
HiPSC-derived bronchial airway epithelium at 45 days of differentiation (iALI) (A) Epithelial cells: Optical microscopy image (left panel), and E-cadherin expression (right panel). iCOPD9 cell line (B, C) Basal cells: TP63 and KRT5 expression, Hy03 and iCOPD9 cell line, respectively. (D) Multi-ciliated cells: expression of the terminal differentiation marker TUBIV, iCOPD2 cell line. (E) Rare clusters of CGHA-positive neuroendocrine cells, iCODP9 cell line. (F) Muc5AC-positive goblet cells, iCOPD9 cell line. (G) Immunostaining of CSSP+ club cells and MUC5AC+ goblet cells in cultures grown without DAPT. Note the presence of CCSP/MUC5AC double-positive cells, iCODP9 cell line. (H) Dot blot analysis to detect the presence of MUC5AC in supernatants of one iALI bronchial epithelium culture (derived from Hy03 cell line) from Day 28 to Day 44. (I) Alcian blue staining and Periodic acid Schiff (PAS), labelling mucus in supernatants of iCOPD9 culture. (J) CCSP quantification at Day 45 in supernatants from iALI bronchial epithelium cultures derived from the HY03, iCOPD9, and iCOPD8 hiPSC lines (K) Quantitative PCR analysis to assess the expression of Foxj1 (ciliated cells), SFTPB (alveolar cells), CHGA (neuroendocrine cells) and Muc5AC (goblet cells). ALI airway epithelium control was obtained from a bronchial biopsy cultured in Lonza BEGM culture medium Panel Scale bar: 20μm

#### iALI generates both major and rare solitary human airway epithelial cells

After 45 days of differentiation, the main bronchial epithelium cell types were observed: basal cells (KRT5 and TP63), ciliated cells (tubulin beta 4, TUBIV), goblet cells (mucin-5AC, MUC5AC), club cells (CCSP, SCGB1A1) and neuroendocrine cells (chromogranin A, CHGA) (figure 5B to F). Club cells and goblet cells could be detected in iALI culture as early as day 14 (figure 5G and I). MUC5AC positive cells were detected by immunofluorescence (figure 5E and G) and supported by a protein release in the supernatant detected by Dot blot analysis, alcian blue staining and Periodic acid Schiff (PAS) (figure 5H and I). Interestingly, we were able to detect either CCSP^+^/MUC5AC^−^ cells and CCSP^−^/MUC5AC^+^ but also a small number of double positive CCSP^+^/MUC5AC^+^ cells as soon as day 14 post ALI (figure 5G), confirmed by colocalization confocal analysis (Supplemental figure 5C). Scanning electron microscopy (SEM) revealed the formation of mucin bundles in culture (Supplemental figure 5A, right lower panels). Concentration of secreted CCSP ranged from 23 to 486 ng/mL, depending on the cell line and experiment (figure 5H). Neuroendocrine cells were also detected both at mRNA and protein level, assessed by CHGA (figures 5F, K). Finally, SEM and TEM acquisitions suggested the presence of another rare epithelial subset of cells harbouring microvilli, also known as brush/tuft cells (Supplemental figure 5A, red asterisk), previously described in proximal airway and in terminal bronchioles [21,22].

#### Functional multiciliated cells airway epithelium

Ciliogenesis was revealed by observation of cilia beating by optical microscopy and by TUBIV immunofluorescent labelling (figure 6A). Multiciliated cells were identified by immunofluorescence labelling only after 21 to 28 days post-ALI in all five iPSC derived lines irrespective of the underlying disease. Dynein axonemal heavy chain 5 (DNAH5) staining was observed throughout the ciliary axoneme (Supplemental figure 5D). The morphology of multiciliated cells was examined using optical microscopy and electron microscopy, either by SEM or TEM (figure 6B-C). TEM cilia structure was characterized as expected by a nine peripheral doublet and a central pair of singlet microtubules (figure 6C), specific of motile cilia [23].

**Figure 6.**
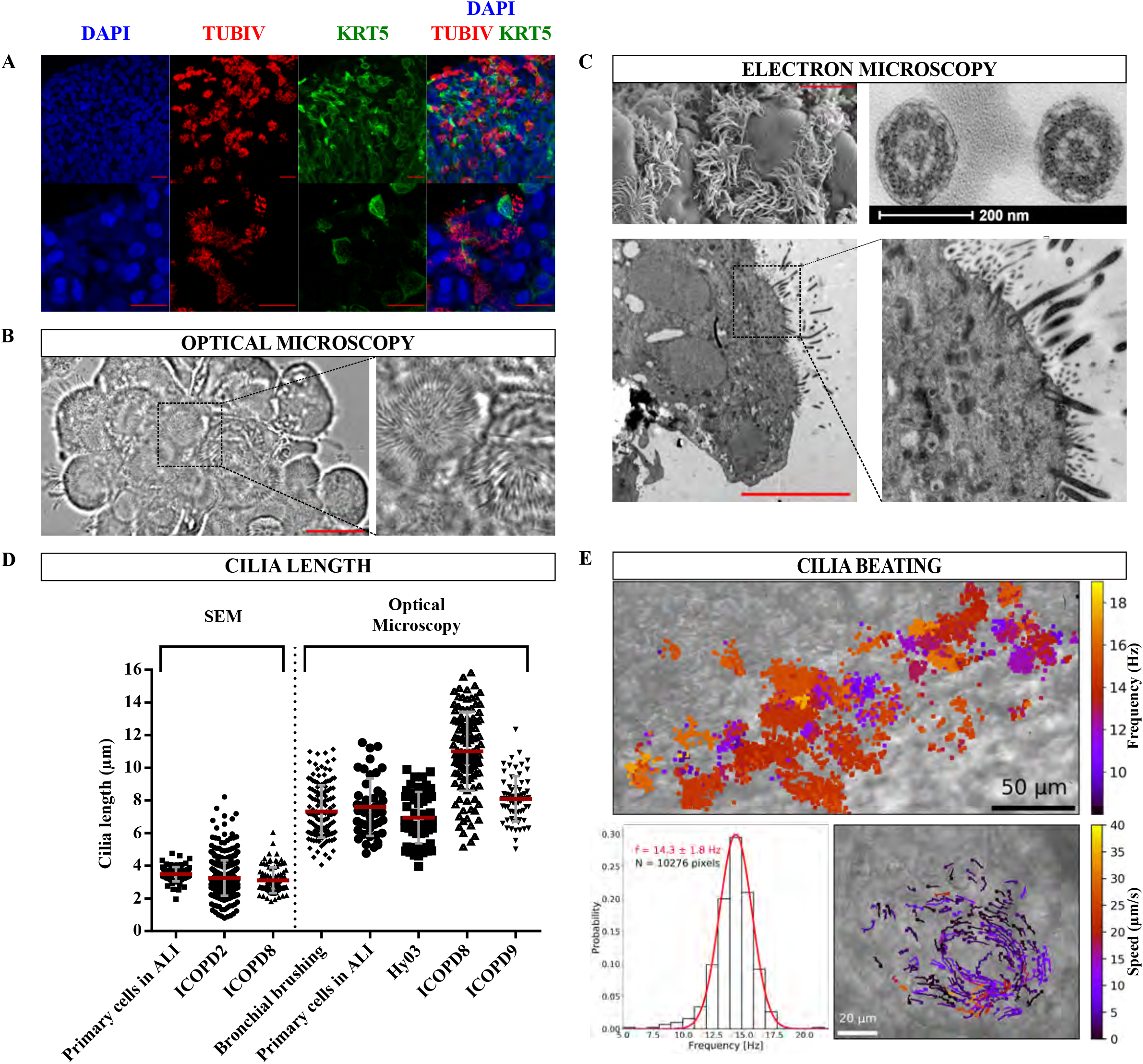
Multi-ciliated bronchial epithelium and cilia characterization at 45 days of differentiation. (A) Confocal microscopy analysis of TUBIV (ciliated cell marker) and KRT5 (basal cell marker) expression, iCOPD2cell line. (B) Optical microscopy images of ciliated cells used for cilia length determination, Hy03 cell line. (C) Top left: Scanning electron microscopy (SEM) image of ciliated cells used for cilia length determination. iCOPD9 cell line. Scale bar 10μm. Right panels: Cilia cross sections by transmission electron microscopy. (D) Determination of cilia lengths by SEM and optical microscopy according to cell lines. Cilia length measurement was performed respectively on primary cells in ALI, n=91 by SEM and n=45 by O.M, bronchial brushing from COPD patients n=141 by O.M, iCOPD2 n=428 by SEM, iCOPD8 n= 98 by SEM and n=120 by O.M, HY03 n=51 by O.M, iCOPD9 n=66 by O.M. (E) Top: Ciliary beating frequency map from a movie (500 frames per second), iCOPD2 cell line Scale bar 50μm. Bottom left: Mean ciliary beating frequency distribution. Bottom right: vectors representing the orientation and celerity of the vortex flow generated by ciliary beating, iCOPD8 cell line Scale bar 20μm. Hz=hertz

Cilia length in iALI was assessed by both optical microscopy and SEM and compared with freshly acquired epithelial cells obtained during endoscopic brushing and classical ALI-cultured airway epithelium. Average cilia length was roughly similar in ALI and iALI when measured either by optical microscopy or SEM (figure 6D). No obvious difference in cilia length was observed between COPD patient-derived iALI and healthy patient-derived iALI or ALI or bronchial brushing. We were able to observe cilia beating using a high-speed camera after isolation of patches of iALI epithelium (Supplemental movie 1), but also on Transwell membrane (Supplemental movie 2). In addition, we acquired cilia beating by live immunostaining using SiR-conjugated fluorogenic probes, SiR-tubulin (Supplemental movie 3).

To establish the mucociliary clearance capacity of the 2D cultures, CBF and mucociliary flow were recorded. iALI cultures had a CBF of 14.3±1.8 Hz, consistent with the frequency of ciliated cells from ALI-cultured primary airway epithelium (figure 6E) [24].

Cultures presented structures with high density of ciliated cells actively beating, giving rise occasionally to localized vortexes (figure 6E, left bottom panel, Supplemental movie 2). The estimated flow velocity of the vortex was approximately 5.6±6.5 μm/s. The iALI bronchial epithelia maintained beating cilia for more than 300 days without cell passaging and without aneuploidy appearance (Supplemental figure 2B). Moreover, cultures could be passaged at least three times after iALI generation.

## Discussion

In this study, we described the generation of iALI bronchial epithelium highlighting an attractive alternative to animal models and *ex vivo* cultures of differentiated bronchial epithelium from endobronchial biopsies. Our differentiation protocol offers a virtually unlimited source of homogeneous reliable human bronchial epithelium. Importantly, this protocol was carried out successfully by four different members of the research group, at least 3 times for each cell lines.

We identified several critical factors that ensure the efficiency and reproducibility of airway epithelium differentiation from human PSCs. First, reprogramming and differentiation were achieved in the same facility by the same team and we do believe this greatly helped for optimally handling hiPSC and decide when to optimally start differentiation for example, selection of clones should be cautious, relying on 1) absence of peripheral signs of spontaneous differentiation, 2) differentiation abilities assessed by levels of CXCR4 expression at the DE stage and 3) ruling out genetic abnormalities by karyotyping or copy counting approaches [18]. We noted that PSCs must be adapted to single-cell culture to obtain a homogeneous cell seeding. When we tried to plate non-adapted cells as large clumps or at high cell density, cell death was reduced, but differentiation was hampered (figure 2E). This could be explained by sustained expression of pluripotency transcription factors within the clumps and/or by altered YAP/TAZ signalling activity. DE and vAFE cell enrichments (assessed by CXCR4 and NKX2.1 expression) achieved at least in 80% of cells at the relevant step were good predictors of the final success of the differentiation process. Based on the work by Matsuno et al [13], we found that APS induction by activation of the activin A/nodal and WNT pathways for 24h, followed by two additional days of activin A activity and TGFβ pathway inhibition for DE induction, without addition of other cytokines or small molecules during vAFE stage, was the most effective strategy. Both SOX2 and SOX9 were observed at the vAFE stage with double positive cells. These bipotent cells were only found in human in the literature and this increased our confidence in the model as it fits with the progenitor patterning [19]. Another key point was the use of the PneumaCult differentiation medium. This proprietary medium, the composition of which is not disclosed, efficiently promotes the differentiation of primary cells obtained from bronchial biopsies. While we cannot exclude that this medium might contain a NOTCH pathway inhibitor, we nonetheless added DAPT to our differentiation protocol. NOTCH signalling inhibition promotes the differentiation into multi-ciliated cell at the expenses of club cells [25]. This protocol generated epithelia containing double positive CCSP^+^/MUC5AC^+^ cells, single CCSP^+^/MUC5AC^−^ and CCSP^−^/MUC5AC^+^ positive cells but largely predominated by basal and ciliated cells. Interestingly, rare cells such as chromogranin A-expressing neuroendocrine cells and tuft cells were found in our model. Altogether, these features suggest that the generated epithelia reproduced many features of a fully differentiated bronchiolar epithelium [26]. The physiological relevance of the model was reinforced by similar to *in vivo* plugs of mucus evidenced both by Alcian blue and PAS staining, the formation of vortexes of mucociliary clearance, cilia length and CBF matching with physiological data.

Besides its reproducibility and simplicity, our protocol provides a 2D bronchial epithelium, unlike other methods that lead to 3D ciliated organoids [8,10,11]. To the best of our knowledge, these three COPD hiPSC lines are the first described in the literature whereas difficulties could be expected given the relative circulating CD34 deficiency previously reported [27]. Moreover, functional, and genetically stable one iALI derived from a COPD patient could be kept consistently differentiated for nearly 400 days at the time of writing. As expected for a disease with multifactorial genetic susceptibility to environmental triggers (e.g. cigarette smoke), the COPD hiPSC lines used here did not show any obvious differentiation specificities, but more work is needed.

In conclusion, we describe here an easy and reliable method to drive PSC differentiation into 2D multicellular bronchial epithelium. This method is highly reproducible, efficient, does not require a cell sorting step and is achievable from samples of patients with pulmonary polygenic diseases or monogenic diseases.

## Materials and Methods

### Clinical characteristics of reprogrammed and differentiated cells

Patients included were defined as severe, with early-onset COPD as FEV1/FVC less than 0.70 and FEV1 percent predicted less than 50% on postbronchodilator spirometry in subjects less than 55 years of age. Normal donors and patients with primary ciliary dyskinesia (PCD) were recruited in the framework of the CILIPS project. More detailed clinical data are **available online (Supplemental Figure 1, supplemental Tables 1 & 2)**.

### Human ESC and iPSC generation and maintenance

The hiPSC lines PCD02.30 (UHOMi001-A) [16], HY03 (UHOMi002-A), iCOPD2 (UHOMi003-A), iCOPD8 (UHOMi004-A) and iCOPD9 (UHOMi005-A) were reprogrammed using the CytoTune®-iPS 2.0 Sendai Reprogramming Kit (Thermo Fisher Scientific, cat.no A16517), according to the manufacturer’s instructions (unpublished results). Emerging hiPSC clones were mechanically selected and clonally expanded using mechanical passaging at early passages (<10 passages). At least three clones for each patient were maintained and their genetic stability was confirmed (Supplemental figure 2). Pluripotency was confirmed by alkaline phosphatase activity staining, SSEA3/4 and TRA1-60 cell surface expression by flow cytometry as previously published [16]. The human ESC line HD291 was derived in our laboratory [17]. The RSP4 and 131007 iPSC lines were derived by the Safe IPS platform (Montpellier, France) using retroviruses and Sendai vectors, respectively. PSC lines were maintained in undifferentiated state in feeder-free conditions on growth factor-reduced Geltrex (Thermo Fisher Scientific) in E8 medium (Thermo Fisher Scientific). Cells were cultured in 35-mm dishes at 37°C and were dissociated mechanically (under an optical microscope) or into single cells at 90% of confluence (every 4-5 days). Single-cell passaging was performed by adding the Versene solution (Thermo Fisher) at 37°C for 5 min and then seeding at 1:10 to 1:20 ratio with addition of 10μM of the ROCK inhibitor Y-27632 (Tocris). The E8 maintenance medium was changed every day.

### PSC differentiation

Differentiation was carried out as described in figure 2A, using reagents at the concentrations listed in supplemental tables 2 and 3. Cells were plated at high-density (one 35mm dish for two Transwell inserts) on Transwell inserts coated with Geltrex. During the differentiation process, medium was changed every day. Cells were differentiated under hypoxia condition (5% 02, 37°C).

### Statistical analysis

Data are presented as means and standard deviations (s.d. or S.E.M), and graphs were generated with GraphPad (Prism, v 6.01). All shown data are from experiments repeated at least three time. P <0.05 indicated significant differences between groups.

## Supplemental data

**Supplemental Table 1: Baseline characteristics of COPD patients**

COPD = chronic obstructive pulmonary disease. FVC = forced vital capacity. FEV1 = forced expiratory volume, RV= residual volume. GORD = gastro-oesophageal reflux disease, Pa02 = partial pressure of oxygen. WA ratio= Wall Area ratio. Wall thickness was expressed as a ratio of the wall thickness to the total airway diameter (WA ratio) and mean value was calculated for each patient from all the bronchi measured. In this study, quantitatively assessment of emphysema was assessed by the percentage of low attenuation area (LAA%) divided by lung or lobe volume(s). A threshold of - 950 Hounsfield Units (HU) was used.

*: other substance abuse included cannabis, intravenous heroin, Subutex misuse (patient COPD2), and cannabis (patient COPD8).

**: Pulmonary hypertension was diagnosed on transthoracic echocardiography; if abnormal, right heart catheterization was performed.

**Supplemental Table 2: Baseline characteristics of PCD patient**

PCD: Primary ciliary dyskinesia. FVC=forced vital capacity. FEV1=forced expiratory volume.

**Supplemental Table 3: Media composition by culture period**

**Supplemental Table 4: Molecules and used concentration**

**Supplemental Table 5: List of reagents and consumables**

**Supplemental Table 6: List and sequences of the primers used for RT-qPCR**

**Supplemental figure 1. Clinical characteristics of patients.**

(A) COPD patients. Left panel: high-resolution inspiratory CT images showing apical centrilobular, para-septal severe emphysema (column apex). In patients COPD2 and COPD8, bronchiectasis and increased airway wall thickness could also be observed (column base). Right panel: rate of change in forced expiratory volume in 1 second (FEV1) over the years since diagnosis. Loss of lung function (% change from baseline) seems more accelerated in COPD patients in this study, around 20 to 30% during the follow up. The mean rate of FEV1 decline in iCOPD2, iCOPD8 and iCOPD9 was respectively 40 mL/year, 83 mL/year, 65 mL/year. (B) PCD patient. Top-left panel: segregation analysis from the studied family demonstrating recessive inheritance of the CCDC40 mutations. Proband was compound heterozygous for Coiled-Coil Domain Containing 40 (CCDC40) gene, carrying two mutations, [c.1116_1117delCT (Exon 7) and c.3180_+_1G_>_A (Intron 19)]. The parents were found to be carriers, segregation analysis showed that c.1116_1117delCT (Exon 7) mutation was inherited from the father and that the mutation c.3180_+_1G_>_A (Intron 19) was inherited from the mother. Her affected sibling was also carrying both CCDC40 mutations. No consanguinity has been reported in this family. Top-right panel: clinical details for the family are shown in the table. Proband exhibited severe rhinosinusitis affection, exacerbations due to bronchiectasis disease and infertility. Her sibling, died prematurely due to congenital heart disease with heterotaxia, comprising transposition of the great arteries (TGA), single ventricle, pulmonary stenosis and dextrocardia. He also suffered from airway ciliary dysfunction. Lower panel: lung computer tomography (CT) scan of the PCD patient. Bronchiectasis can be identified on inspiratory CT images (a), present mainly in the middle lobe (b). There were thickening of the airway wall (c), central mucous plugs (d) and some area of lung consolidation (e). Small airway disease was illustrated by impaction in bronchioles and small nodules (f); however, expiratory CT image did not demonstrate areas of air trapping (data not shown).

**Supplemental Figure 2: Genetic integrity of the hiPSC lines used for differentiation into iALI bronchial epithelia**

(A) Genomic integrity evaluation of the hiPSC lines using the iCS-digital test [18]. Copy number variation analysis using droplet digital PCR and DNA extracted from the different hiPSC lines in culture (iCOPD8, iCOPD9, iCOPD2, PCD02.30 and HY03). All the hiPSC lines remained euploids, except iCOPD2 (clone A13) that displayed a copy number gain on chromosome 20q at mechanical passage 70 and clumps passage 3 (M70CL3) and was therefore later on discarded. Error bars indicate the Poisson distribution (95% confidence intervals).

(B) Same analysis, using the iCS-digital Aneuploidy test to screen the 23 chromosomes in one iALI bronchial epithelium culture that was maintained for twelve months in culture.

**Supplemental Figure 3 NKX2.1 and CXCR4 FACS gating strategy**

**(A)** Gating strategy for the isolation of CXCR4 positive cells.

Flow cytometry gating strategy to viable CXCR4 cell subsets at the Definitive endoderm stage. Staining of single-cell solutions using isotype and CXCR4 conjugated antibodies and analysis by flow cytometry. (A-C) Gating strategy to exclude doublet (B) and isolate real single-cell unit (C). Among single cells, live cells were selected on absence of Zombie violet staining (D). (E) CXCR4 expression of cells compared to isotype control on PE staining **(B)** Gating strategy for the isolation of NKX2.1 positive cells.

(A) Flow cytometry gating strategy to viable NKX2.1 cell subset at the Anterior Foregut Endoderm stage. Staining of single-cell solutions using different unconjugated antibody and analysis by flow cytometry. (A-C) Gating strategy to exclude doublet and isolate real single-cell unit. (D) Among single cells, live cells were selected on absence of Zombie violet staining (E) NKX2.1 expression of cells compared to isotype control on Alexa 488 staining.

**Supplemental Figure 4. Characterization of v-AFE progenitors SOX9 expression and time course expression of NKX2.1 during v-AFE induction.**

(A) Kinetic expression of NKX2.1 by immunostaining during v-AFE induction, in Hy03 cell line. Note increasing expression during the time course. Scale bar: 20μm.

(B) Immunostaining of Hy03 -derived v-AFE cells for SOX9 (green) nuclear proteins. during v-AFE stage. Scale bars: 20 μm.

(C) Immunolabelling of Hy03 cell line v-AFE stage for NKX2.1 (orange) and ki67 (green). Nuclei counterstained with DAPI.

(D) Immunofluorescence of v-aFE stage showing no contamination by AFP positive (liver) cells (PCD02.30 cell line). These results were available for all the others iPSC cell lines (data not shown). HepG2 hepatoma cell line was used as positive control, characterized by AFP expression (red). Scale bar 50μm.

**Supplemental Figure 5: iALI**

(A) Electron microscopy of hiPSC derived airway epithelium grown at an air–liquid interface (iALI) after 45 days of differentiation (iCOPD2 and iCOPD9 cell lines).

Column 1: Transmission Electron Microscopy of mature epithelium. Top: presence of two contiguous ciliated cells. Red arrows head show epithelial features highlighted by tight junction and desmosomes. Bottom: at the apical pole, red stars parts of cilia section.

Column 2: Scanning electron microscopy of epithelial layer. Top: goblet cell layer; middle: red arrows indicate cilia of multiciliated cell and orange ones indicate mucus globules. Bottom: red asterisks show microvilli. Orange arrow indicate cluster of mucus. Red arrow exhibit multiciliated cells. Scale bar is 10 μm.

(B) Tight-junction integrity during the course of culturing was assessed by measuring the TEER. At least three inserts were analysed at each point of the time course and the data represents the mean +/− SD (iCOPD9 cell line).

(C) Colocalization of two colours confocal image of CCSP/MUC5AC. MUC5AC protein (red) colocalizes with CCSP protein (green) at day 14 of ALI. Orthogonal views (XY, XZ, YZ) illustrating colocalization of CCSP and MUC5AC. Colocalization is visible as a yellow colour. iCDOP9 cell line. Scale bar is 20 μm.

(D) Multiciliated cell characterization using DNAH5 and TUBIV antibody. DNAH5 immunolabelling exhibit an axoneme in iCOPD9 cell line (left panel). TUBIV immunofluorescence show only a cilia staining (middle). Merge of DNAH5 and TUBIV staining (right panel), iCODP9 cell line. Scale bar is 10 μm.

**Supplemental Movie 1: iALI bronchial epithelium obtained from the iCOP9 hiPSC cell line (X20)**

Cilia beating was clearly visible on this video from iCOPD9 at Day 45. Acquisition was performed with inverted microscopy during cell culture routine check-up. The video represents cells that have been dissociated, before passaging cells into another coated Transwell. Dissociation was performed using Trypsin 10 minutes at 37°.

**Supplemental Movie 2: iALI bronchial epithelium obtained from the iCOPD8 hiPSC cell line (X40)**

This acquisition was used for biophysics analysis i.e. Ciliary Beat Frequency (CBF). The video was recorded at 500 frames per second. This video includes 1500 frames.

**Supplemental Movie 3: iALI bronchial epithelium obtained from the iCOPD9 hiPSC cell line and with live immunofluorescence for TubIV**

**Supplemental Information: Supplemental Methods and Supplemental Clinical data**

## Acknowledgements

We are grateful to Pascal Chanez and Delphine Gras for reagents. This work was supported by the grants FDM20170638083 (FRM) and RF20160501664 (Vaincre la Mucoviscidose), and by Boehringer Ingelheim. The clinical studies were conducted by DRCI CHU de Montpellier with protocol numbers UF 9791 and UF 9174.

We wish to thank Montpellier Rio Imaging for access to their imaging technological facilities (http://www.mri.cnrs.fr/en/). We thank Elena Hauser for her technical help. We thank Dr Sebastien Bommart for chest CT scan analyses. We thank iPSC platform for RSP4 and 131 000 iPSCs cell lines.

## Author contributions

E.A., M.F., S.A., A.B., and J.D.V designed the study and analyzed data; E.A., M.F., S.A., C.B., J.M., A.P., C.V., M.J., C.C., H.B. performed the experiments, collected and analysed data; M.J. and G.M. analysed biophysical data .E.A., M.F., S.A., I.V., J.P.G, A.B., and J.D.V wrote the paper. All authors approved the final version prior to submission. All authors have read, reviewed and approved the final submitted manuscript and agree to take public responsibility for it.

