## Supplemental Information for "Differentiation of human induced pluripotent stem cells into functional airway epithelium"

Includes supplemental clinical data and supplemental methods.

**Supplemental Clinical Data**

### Supplemental information of PCD and COPD history

**COPD patient’s characteristics and history**

Patients with severe COPD were recruited in the framework of the INVECCO project. Patients exhibited normal blood level of α-1 anti-trypsin and none of them carried any TERT mutations, to avoid any monogenic form of COPD.

Early onset of disease was suggested by early onset of symptoms, at an average age of 35 years.

Disease was revealed by dyspnea, pneumothorax or acute exacerbation. The average time between COPD diagnosis and first symptoms was 12 years. Familial history of COPD at first degree was found in 2 out of the 3 patients. All of them were at least GOLD stage 3. Environmental exposures were characterized by early and heavy active tobacco exposure, and all of them had in utero exposure to tobacco and second-hand smoke in early childhood. The average smoking history was 50 pack-years (range, 30–75). Others toxic consumption such as cannabis and parenteral heroin were also reported, in agreement with the literature that already reported an association with severe emphysema ^16^. The dyspnea was severe, requiring long term home supplemental oxygen support for 2 patients. No cardiovascular, diabetes, cancer or pulmonary hypertension comorbidities was described, but they usually had severe osteoporosis despite their young age. 2 of 3 COPD patients had spontaneous pneumothorax, recurrent for both. Notably, 2 of 3 individuals suffered of three or more exacerbations per year requiring hospitalization. All three patients received at baseline long-acting beta2-agonist (LABA) and anti-cholinergic (LAMA) drugs but not inhaled corticosteroid (ICS). Clinical course is detailed in Supplemental information for the three patients. Chest CT scans showed severe apical centrilobular emphysema destruction, basal bronchiectasis and increased wall thickness (figure 1A, left panel). Lung function fast declined in these COPD patients. Mean level of change in FEV1 (ml/yr.) was −25.3 (SD, 43.3) ml/year (figure 1A, right panel).

Diagnosis was established on respiratory symptoms, in the context of severe sibling PCD. She presented mild lung phenotype disease with normal lung function at baseline, with less than 1 exacerbation per year, rhinosinusitis, and infertility (Table 2). Segregation analysis and clinical features in the family is detailed in figure 1B, top panel. CT scan mainly showed bronchiectasis and mucous plug impactions (figure 1B, bottom panel).

At the time of the inclusion, all the patients of the COPD group were on a lung transplantation waiting list. At the time of the manuscript submission, iCOPD2 patient was admitted in intensive care unit for a severe COPD exacerbation requiring mechanical ventilation. iCOPD8 patient declined lung transplantation. iCOPD9 patient was programmed to undergo single lung transplantation (pre-transplant pleurodesis due to iterative pneumothorax). The lung transplant project was stalled because of overweight for more than 24 months. Unfortunately, a localized lung adenocarcinoma was recently discovered. The patient will be proposed to undergo bronchoscopic lung volume reduction due the severe emphysema phenotype.

**PCD patient clinical characteristics and history with disease-causing variants in CCDC40**

The healthy control aged 41 at inclusion had normal lung function and suffered no respiratory symptoms from childhood and had no familial history of any chronic airway disease. PCD patient was a non-smoker female 34 years old at inclusion (Table 2). Disease was associated with bi allelic mutations in Coiled-Coil Domain Containing 40 (CCDC40) gene [c.1116_1117delCT (Exon 7) and c.3180 + 1G > A (Intron 19)] (figure 1B, top panel). Diagnosis was established on respiratory symptoms, in the context of severe sibling PCD. She presented mild lung phenotype disease with normal lung function at baseline, with less than 1 exacerbation per year, rhinosinusitis, and infertility (Table 2). Segregation analysis and clinical features in the family is detailed in figure 1B, top panel. CT scan mainly showed bronchiectasis and mucous plug impactions (figure 1B, bottom panel). Clinical course is described in Supplemental information.

Proband was included in this study at the age of 34 years old. She presented symptoms since the childhood, mainly represented by sinusitis and lung exacerbations. The annual rate of exacerbations was lower than 2/year, and no one lead to hospitalization. Decline of lung function was observed during the follow up, from normal lung volumes to a decrease of 50% of FEV1 at the time of manuscript submission. She also suffers from infertility, no ectopic pregnancy neither miscarriage occurred. Successful pregnancy was achieved by in vitro fertilization with oocyte donation.

**Supplemental Methods**

### Live imaging

Cells were incubated in culture medium containing 1µM SiR-tubulin and verapamil (SiR-tubulin kit, Spirochrome) at 37°C for 60 min. Live images were acquired using a two-photon microscope LSM 7 MP OPO (Zeiss, France) with an upright Axio Examiner Z.1 optical microscope associated with a femtosecond Ti: sapphire laser (680–1080 nm,80 MHz, 140 fs, Chameleon Ultra II, Coherent, France) pumping a tuneable OPOs (1000–1500 nm, 80 MHz, 200 fs, Chameleon Compact OPO, Coherent, France) (Software Zen Ver. 2012). A x20 water immersion objective (W Plan Apochromat DIC VIS-IR) was used for time-lapse image acquisition with the following characteristics: 300x300 pixels (141,7 x141,7 µm) frame size, scan speed (Pixel Time 1.34 μs, Frame Time 0.14 s), time-lapse of 16.55 s (55 frames), OPO excitation wavelength of 1097 nm, and narrow band pass filter at 650-705 nm in front of one of the detectors to detect the fluorescence.

### Determination of Cilia Beating Frequency (CBF) and flow velocity

In each experiment, more than 50 ciliated cells corresponding to more than 10 000 pixels/ movie and 100 trajectories in each hiPSC line, were analysed for calculating the CBF and flow velocity, respectively.

**CBF**

All the samples were observed immediately after taking out of the incubators to keep a condition of 5% CO2 at 37°C. CBF were determined for Hy03, iCOPD2, 8 & 9 cell lines, for at least 2 independently experiments.

The mean ciliary beat frequency was determined from videos obtained with an inverted optical microscope in bright field (Leica DMI 3000 B), equipped with a x63 objective (N.A. 0.7) and with a high speed camera (Photron Fastcam PCI 1024), at 37°C and 5% CO2. Ciliary motions were recorded at 500 fps for 3 seconds. Analysis of CBF was processed on 280µm square region of interest. The periodic variation in intensity of pixels located on the path of beating cilia are used to infer the beating frequency. For each pixel, the power spectrum (squared FFT (Fast Fourier Transform) versus frequency) shows a peak (maximum occurring periodicity) which is taken as the beating frequency. For each recording, the frequencies are mapped, and their histogram is fitted by a Gaussian distribution to determine the mean beating frequency and its standard deviation.

**Mucous flow velocity**

Dead cells embedded in the mucus are tracked and their trajectories are used to describe the mucus flow. Movies were obtained under a Leica microscope with an objective x40 or x63, and using a high-speed camera (Photron Fastcam PCI 1024) at 60 or 125 fps for a total of 1536 frames per recordings (12 s at 125 fps or 25 s at 60 fps).

Dead cells tracking was performed using the trackpy library in Python (version 0.3.2; Allan, 2016 DOI [10.5281/zenodo.60550](https://www.researchgate.net/deref/http%3A%2F%2Fdx.doi.org%2F10.5281%2Fzenodo.60550)). For each tracked dead cell, velocities computed on 10-frames-long portions of its trajectory are represented in a histogram. The velocity corresponding to the maximum of this histogram is taken as the estimated dead cell velocity. Fig.7E is a map of the dead cells trajectories, the colour code corresponding to the estimated flow velocity.

Estimated flow velocity of dead cells was approximately 2.2 +/- 4.3 µm/s in iCOPD8 hiPSC derived airway epithelium and 38 ± 28 µm/s in iCOPD9. Experiments were performed using hiPSC (iCOPD8, iCOPD9 cell lines) on 3 wells per cell line. Normal human bronchial epithelial cell (NHBEC) were used as control. Normal human bronchial epithelial cell (NHBEC) were used as control. Range of mucous flow velocity of HBECs was between [0-50 µm/s].

### Genomic stability

Genomic stability was assessed by detection of recurrent genetic abnormalities in hPSCs using the droplet digital PCR technology, provided as a service by Stem Genomics, as described previously ^18^.

### Cytospin preparation

Cells were dissociated to single cells to reach a density of 500 000 cells/ml. 60 µl of cell suspension was dropped on cytospin slides that were centrifuged at 500 rpm for 5 min. Then, cells were fixed in cold acetone at -20°C for 10 min. Excess acetone was removed, and dry slides were stored at 4°C.

### Immunofluorescence and phosphatase alkaline activity

Samples were fixed in 4% paraformaldehyde (PFA) at room temperature (RT) for 15 min, and then permeabilized with PBS containing 0.5% Triton X-100 for 15 min. Cells were blocked with 10% of donkey serum in PBS containing 1% BSA and 0.1% Triton X-100 at RT for 60 min. Primary and then secondary antibodies were diluted in 1% BSA/0.1% Triton X-100/PBS. Samples were incubated with primary antibodies at 4°C overnight and with secondary antibodies at RT in the dark for 60 min. Nuclei were stained with DAPI (1:5000) for 3 min and then samples were mounted in ProLong Gold (Thermo Fisher) and stored in the dark at RT. A Zeiss microscope (LMS700) was used for image acquisition and data were analysed with ImageJ (v1.52i).

Phosphatase alkaline activity was assessed using the ScienCell Research Laboratories (#8288) kit following the manufacturer’s instructions.

**Scanning Electron Microscopy (SEM)**

Transwell inserts were fixed with 2.5% glutaraldehyde in PHEM buffer (pH 7.2) at RT for 60 min, followed by washes in PHEM buffer. Fixed inserts were dehydrated in increasing concentrations of ethanol, from 30 to 100%. Samples were immersed in ethanol – hexamethyldisilazane (HMDS) solution for 10 min and then in HMDS alone. Transwell inserts were sputter coated with a 10nm-thick gold film and analysed with a scanning electron microscope (Hitachi S4000; MRI facility, INM Montpellier France) using a lens detector with an acceleration voltage of 10KV at calibrated magnifications.

### Transmission Electron Microscopy (TEM)

Transwell inserts were fixed in 2.5% glutaraldehyde/PHEM buffer (1X, pH 7.4) at 4°C overnight. Samples were rinsed with PHEM buffer and post-fixed in 0.5% osmic acid/0.8% potassium ferrocyanide in the dark at RT for 2h. Inserts were washed in PHEM buffer twice, and then dehydrated in a graded series of ethanol solutions (from 30 to 100%). Samples were embedded in EmBed 812 using an Automated Microwave Tissue Processor for Electronic Microscopy (Leica EM AMW). Thin sections (70 nm; Leica-Reichert Ultracut E) were collected at different points. Sections were then stained with 1.5% uranyl acetate/70% ethanol/lead citrate and observed using a Tecnai F20 transmission electron microscope at 120KV (MRI facility, INM Montpellier France).

### Cilia length determination

Cilia length was measured using the line measurement tool in ImageJ on SEM or Optical microscopic images. To limit measurement bias, only whole flat cilia parallel to the plane of focus were analysed. Angled cilia or cilia with incomplete visible part were excluded. Cilia length was assessed by the average of three measurements ^1^. At least 20 ciliated cells were measured to determine the average ciliary length.

### Flow cytometry analysis

For extracellular CXCR4 staining, cells were first incubated with Zombie violet (1:1000, Biolegend) to differentiate between live and dead cells at RT in the dark for 15 min. Cell pellets were then incubated with an anti-CXCR4 conjugated to PE antibody (Mouse PE, 1:200, BD Biosciences) or its isotype control at 4°C for 30 min.

For intracellular NKX2.1 staining, after staining with Zombie violet to assess cell viability, cells were fixed in 0.5% PFA/PBS at RT for 10 min. Cells were blocked and permeabilized by incubation with 1X saponin (10X, Thermo Fisher) and 10% donkey serum (Sigma) in MilliQ water at RT for 20 min. Then, cells were incubated with unconjugated anti-NKX2.1 primary antibody (rabbit, 1:1000, Abcam) at RT for 30min followed by an AlexaFluor 488 secondary antibody (donkey anti-rabbit, 1:3000, Life technologies) at RT in the dark for 60min. Antibodies were diluted in MilliQ water/1X saponin. All acquisitions were done on a Beckman Coulter Gallios cytometer and data were analysed with the Kaluza software. The detailed cytometer gating strategy is described in Supplemental figure S3.

### RT-qPCR analysis

Reverse transcription (RT) was performed using SuperScript™ First-Strand Synthesis System (ref 11904-018, Invitrogen) as recommended by the manufacturer in 20 μL reaction volume that included 1 µg RNA (extracted with RNeasy Mini Kit, ref 74106, Qiagen), Superscript II RT, oligo-dT primer, dNTP mixture, MgCl_2_, 0.1 M DTT and RNase inhibitor. Quantitative PCR (qPCR) was then performed using the LightCycler® 480 SYBR Green I Master (04707516001, Roche) with 2 μl of the RT reaction product (1:20 dilution) and 0.5µl of each primer diluted at 10µM (Integrated DNA Technologies IDT) in a total volume of 10 μl. PCR amplifications were carried out using a LightCycler 480 apparatus and the following programme: 1 cycle of 95°C for 10s; 40 cycles of 95°C for 10s, 60°C for 15s; 72°C for 15s, 1 cycle 95°C for 5s, 65°C for 1min, 97°C forever. Gene expression levels were normalised to the expression of the housekeeping gene *GAPDH*, using the following formula: 2^-ΔΔCt^, where ΔΔCt = ΔCt unknown - ΔCt positive control. Each sample was analysed in duplicate and multiple controls were included. Primary Human Bronchial Epithelial Cells (HBECs) cultivated in ALI culture conditions were used as lung positive control. Contamination from other layers was assessed using control samples: liver (HepG2 cells), thyroid (Thermo Fisher Scientific‎, cat no. QS0631), brain (Thermo Fisher Scientific‎, cat no. QS0611), and colon (Thermo Fisher Scientific‎, cat no. QS0613) total RNA. Primer sequences are shown in Supplementary table 4.

### Quantification of SGB1A1 and MUC5AC secretion

Apical secretions were gently collected with a micropipette and stored at -80°C°C, for a maximum of one month. Typically, to collect the secretions for one culture transwell, we performed an apical wash with 200µL of PBS warmed at room temperature. One up and down pipetting was performed to collect PBS, that also allows some mucus to detach. SGB1A1 and MUC5AC were quantified from these samples.

SCGB1A1 and MUC5AC secretions were measured in a dot blot assay using primary antibodies (anti Muc5Ac (45M1); ThermoFisher Scientific or Club Cell Protein rabbit polyclonal antibody ; Biovendor). Apical lavages were spotted onto a nitrocellulose membrane. The membrane was then incubated in Odyssey blocking solution. After washing, the membrane was incubated with primary antibodies at room temperature for 2 hours. Secondary fluorescent anti-mouse and anti-rabbit antibodies was then added to the membrane. Positive signals were detected using an Odyssey imager. Optical densities were measured with Alphaview software.

For MUC5AC detection: apical lavages from Day 28 to Day 46 of one iALI bronchial epithelium culture (derived from Hy03, iCOPD8 and iCOPD9) were spotted onto a nitrocellulose membrane. Negative controls (PBS and ALI medium) and positive controls (supernatants of ALI culture bronchial epithelium from biopsies) were added to check the specificity of the experiment. Supernatant of hiPSC at Day 0 was spotted to confirm the lack of MUC5AC secretion.

For SCGB1A1 detection: supernatants of iALI bronchial epithelium cultures derived from the HY03, iCOPD9, and iCOPD8 hiPSC cell lines were spotted. Negative control (PBS) and positive control (supernatant of ALI culture bronchial epithelium from biopsies) were added to check the specificity of the experiment. 0.625 to 5ng of Recombinant human Club Cell protein (Biovendor) were used as standard range to quantify SCGB1A1 secretion.

### TEER measurement

TEER: Epithelial monolayer integrity was assessed by trans-epithelial electrical resistance (TEER) using WPI EVOM2 Model with STX2 electrode. Measurements were done on transwell devices with 1ml of PneumaCult-ALI Medium in the basolateral chamber and 300µl of PBS calcium, magnesium (Gibco, cat no 14040083) at the top. The measurement process consists on measuring the blank TEER (TEER_BLANK_) of the semipermeable membrane only coated with Geltrex and without cells. Total TEER (TEER_TOTAL_) was a measure of the resistance across the cell layer on the insert semipermeable membrane. The real TEER of the epithelial cell was calculated with the following formula TEER cell layer (Ω) = TEER_TOTAL_ – TEER_BLANK_. TEER cell layer (Ω) was then multiplicated per the area of the insert (1.12cm²) to obtain final TEER in Ω.cm².

1.	Dummer, A., Poelma, C., DeRuiter, M. C., Goumans, M. J. & Hierck, B. P. Measuring the primary cilium length: improved method for unbiased high-throughput analysis. Cilia 5, 7 (2016).
