## Supplemental Table 1 for "Differentiation of human induced pluripotent stem cells into functional airway epithelium"

|  | **COPD2** | **COPD8** | **COPD9** |
| --- | --- | --- | --- |
| Age (years) | 52 | 46 | 54 |
| Sex | F | M | M |
| Body-mass index (kg/m2) | 30 | 24 | 32 |
| **Smoking**  Current/ Ex-smoker  Pack-year history  Age of beginning (years)  **Others toxic *** | Current  45  14  Yes | Ex-smoker  30  18  Yes | Ex-smoker  75  24  No |
| **Pulmonary function and symptoms**  Age of diagnosis  Age at onset of first symptoms (years)  In utero smoking exposure  Premature birth  Familial history of CODP/emphysema  Increased dyspnea  Increased cough  Medical Research Council dyspnea scale | 48  40  Yes  No  Yes  Yes  Yes  4 | 43  30  Yes  No  No  Yes  No  1 | 49  35  Yes  Yes  Yes  Yes  Yes  4 |
| **Lung function**  FEV (L)  FEV (%)  FEV1/FVC ratio (%)  RV (L)  RV (%)  DLCO (%) | 0.35  14  32  4.39  244  30 | 1.19  34  25  3.21  156  41 | 0.97  30  36  4.72  213  35 |
| **Medication**  Long acting β2 agonist  Long-acting muscarinic antagonist  Inhaled corticosteroid  Use of benzodiazepine  Azithromycin | Yes  Yes  No  Yes  Yes | Yes  Yes  No  Yes  No | Yes  Yes  No  Yes  No |
| **Clinical findings**  Oxygen supply (L/min)  BODE index  Exacerbation rate in previous 12 months  Pneumothorax  Osteoporosis  Pulmonary hypertension******  Coronary Artery Disease  Lung cancer  Type 2 diabetes  GORD  Anxiety/Depression | 1  6  3  No  Yes  No  No  No  No  Yes  Yes/No | 0  3  3  Yes  No  No  No  No  No  Yes  Yes/Yes | 3  6  1  Yes  Yes  No  No  No  No  Yes  Yes/Yes |
| **Blood test**  Eosinophils (10^9 cells per L)  Eosinophils (% total WBC)  α1 anti trypsin (g/l)  Pa02 (mmHg, ambiant air)  PaC02 (mmHg, ambiant air) | 150  2.4  1.16  62  48.8 | N.A  N.A  1.37  82  32.6 | 810  5.9  1.1  76.6  44.4 |
| **CT scan**  Emphysema (wall lung) %  WA ratio mean (min-max) | 18.4  59.75 (48.1-68.1) | 12.99  57 (44.9-69) | 31.95  68.2 (57.7-77.1) |
