## Supplemental Table 2 for "Differentiation of human induced pluripotent stem cells into functional airway epithelium"

|  | **PCD patient** |
| --- | --- |
| Age (years) | 30 |
| Sex | F |
| Smoking | No |
| **Baseline medical history**  Age at onset of first symptoms (years)  Familial history of pulmonary disease  Chronic productive cough  Medical Research Council dyspnea scale  Fertility problems  Situs Inversus  Chronic Otitis Media  Rate of exacerbations (per year)  PCD-related hospitalizations  Chronic nose symptoms  Retinitis pigmentosa  Kidney problems  Gastrointestinal problems  Surgical interventions related to PCD  Non-PCD-related comorbidities | > 2 years  Yes  Yes  2  Yes  No  No  1  No  Yes  No  No  No  No  No |
| **Diagnosis**  Nasal NO  Genetic analysis | N.A  Heterozygous mutations CCD40 gene |
| **Lung function**  FEV (L)  FEV (%)  FEV1/FVC ratio (%)  DLCO (%)  Pa02 (mmHg) | 1,91  69  75  80  72 |
| **Medication**  Long acting β2 agonist  Long-acting muscarinic antagonist  Inhaled corticosteroid  Azithromycin  Physiotherapy  Oxygen supplementation | Yes  No  No  No  Yes  No |
| **Imaging findings**  Bronchiectasis  Atelectasis  Infiltrations  Lobar collapse  Mucus plugging  Location of bronchiectasis  Pansinusitis | Yes  No  Yes  No  Yes  Middle lobe  Yes |
| **Microbiology** | No chronic colonization |
