## Supplemental Table 3 for "Differentiation of human induced pluripotent stem cells into functional airway epithelium"

| Media | Induced Pluripotent Stem cells | Anterior Primitive Streak | Definitive Endoderm | Anterior Foregut Endoderm | Lung  Progenitor | Lung Airway Epithelium |
| --- | --- | --- | --- | --- | --- | --- |
| Days | Day 0 | Day 1 | Day 2-3 | Day 4-8 | Day 9-10 | Day 11-42+ |
| Basal media | Essential 8 Medium Penicillin-Streptomycin (10,000 U/mL) 1% | RPMI 1640  1x B27 supplement (50x), minus vitamin A  Penicillin-Streptomycin (10,000 U/mL) 1% | RPMI 1640  1x B27 supplement (50x), minus vitamin A  Penicillin-Streptomycin (10,000 U/mL) 1% | RPMI 1640  1x B27 supplement (50x), minus vitamin A  Penicillin-Streptomycin (10,000 U/mL) 1% | PneumaCult-Ex Plus  Medium*  Penicillin-Streptomycin (10,000 U/mL) 1% | PneumaCult-ALI  Medium**  Penicillin-Streptomycin (10,000 U/mL) 1% |
| [Add](https://www.linguee.fr/anglais-francais/traduction/extemporaneous+preparation.html) the day of use | + 10µM Y-27632 | + 10µM Y-27632  + Activin A 100ng/ml  + CHIR99021 3µM | + 10µM Y-27632  + Activin A 100ng/ml  + LDN-193189 250nM | No cytokines | No cytokines | 10 µM DAPT from day 28 to day 42 |
