## Supplemental Table 4 for "Differentiation of human induced pluripotent stem cells into functional airway epithelium"

| Growth  Factor | Stock  Concentration | Final  Concentration |
| --- | --- | --- |
| Activin A | 50 µg/ml | 100 ng/ml |
| Y-27632 | 5 mM | 10 µM |
| CHIR99021 | 3 mM | 3 µM |
| LDN-193189 | 450 µM | 250 nM |
| DAPT | 10 mM | 10 µM |
