## Supplemental Table 5 for "Differentiation of human induced pluripotent stem cells into functional airway epithelium"

| **REAGENT** | **SOURCE** | **IDENTIFIER** |
| --- | --- | --- |
| ***Chemicals, Peptides, and Recombinant Proteins*** | | |
| Geltrex LDEV-Free, hESC-Qualified, Reduced Growth Factor Basement Membrane Matrix | Thermo Fisher | A1413301 |
| StemSpan SFEM II | Stemcell | 09605 |
| StemSpan Erythroid Expansion Supplement | Stemcell | 02692 |
| Sodium Butyrate, | Stemcell | 72242 |
| CytoTuneTM −iPS 2.0 Sendai Reprogramming Kit | Thermo Fisher | A16517 |
| B27 supplement (50x), minus vitamin A | Thermo Fisher | 12587010 |
| Y-27632 dihydrochloride | Tocris | 1254 |
| CHIR-99021 | Sigma | SML1046-5MG |
| LDN-193189 | Miltenyi biotec | 130-103-925 |
| DAPT | Tocris | 2634/10 |
| Activin A | Peprotech | AF-120-14E |
| Permeabilization Buffer (10X) | Thermo Fisher | 00-8333-56 |
| Triton X-100 (laboratory grade) | Sigma | X100-5ML |
| Bovine Serum Albumin | Sigma | A7906-100G |
| ProLong Gold Antifade Mountant | Thermo Fisher | P36930 |
| Donkey serum | Sigma | D9663-10ML |
| Goat serum | Sigma | G9023-5ML |
| DAPI | Sigma | D9542-5MG |
| CryoStor CS10 | Stemcell | 07930 |
| Zombie Violet Fixable Viability Kit | BioLegend | 423113 |
| Penicillin-Streptomycin (10,000 U/mL) | Thermo Fisher | 15140122 |
| Versene Solution | Thermo Fisher | 15040033 |
| Glutaraldehyde (25%) | Deltamicroscopies | 16210 |
| Paraformaldehyde (10%) | Deltamicroscopies | 15712 |
| ***Oligonucleotide primers*** | Integrated DNA Technologies (IDT) | See Table S4 - List and sequences of the primers used for RT-qPCR |
| ***RNA controls*** |  |  |
| Human lung | Invitrogen | QS0618 |
| Human thyroid | Invitrogen | QS0631 |
| Human heart | Invitrogen | QS0614 |
| Human colon | Invitrogen | QS0613 |
| Human liver | Invitrogen | QS0617 |
| Human brain | Invitrogen | QS0611 |
| ***Antibodies (dilution)*** |  |  |
| Donkey anti-rabbit IgG AlexaFluor 488 *(1:1000)* | Life technologies | A21206 |
| Donkey anti-mouse IgG AlexaFluor 555 *(1:1000)* | Life technologies | A31570 |
| Donkey anti-goat IgG AlexaFluor 647 *(1:1000)* | Life technologies | A21447 |
| Mouse anti-CXCR4 (PE) *(1:200)* | BD Biosciences | 557145 |
| Mouse IgG2a, (PE) *(1:200)* | BD Biosciences | 556653 |
| Mouse anti-AFP *(1:200)* | Sigma | A8452 |
| Mouse anti-Mucin 5AC *(1:200)* | Abcam | ab3649 |
| Mouse anti-SOX2 *(1:100)* | Abcam | ab79351 |
| Rabbit anti-SOX9 *(1:1000)* | Abcam | ab185230 |
| Mouse anti-β-Tubulin IV *(1:200)* | Sigma | T7941 |
| Rabbit anti-DNAH5 *(1:200)* | Sigma | HPA037470 |
| Rabbit anti E-cadherin *(1:200)* | Santa Cruz Biotechnology | sc-7870 |
| Rabbit anti-Oct 4 *(1:250)* | Santa Cruz Biotechnology | sc-9081 |
| Rabbit anti-NANOG *(1:200)* | Abcam | ab80892 |
| Rabbit anti-CCSP *(1:1000)* | BioVendor | Rd181022220 |
| Rabbit anti-TTF1 (NKX2.1)  *IF: 1:200 / FCM 1:1000* | Abcam | ab76013 |
| Mouse anti-Chromogranin A *(1:200)* | Abcam | ab715 |
| Rabbit anti-Ki67 *(1:200)* | Abcam | ab15580 |
| Goat anti-SOX17 *(1:150)* | R&D Systems | af1924 |
| Goat anti-FOXA2 *(1:150)* | R&D Systems | af2400 |
| SiR-tubulin | Spirochrome AG | SC002 |
| ***Software and Algorithms*** |  |  |
| GraphPad | Prism | Version 6.01 |
| ImageJ | National Institutes of Health | Version 1.52i |
| Inkscape | Free Software Foundation | Version 3 |
| Alphaview sofware | Protein simple | Version 3.4.0.0 |
| Zen 2.3 | Carl Zeiss Microscopy | Version 4.03 |
| Kaluza Analysis | Beckman Coulter | Version 2.1 |
| Kaluza for Gallios Acquisition | Beckman Coulter | Version 1.0 |
| Flow Cytometer (FCM) | Beckman Coulter | Gallios |
| Confocal Microscopy | Zeiss | LMS700 |
| Odyssey imaging system | LI-COR | Odyssey 9120 Model |
| Scanning electron microscopy | Hitachi | S4000 |
| Optical microscopy | Leica | DMi1 Model/MC170HD camera |
| ***Others*** |  |  |
| Essential 8 Medium | Thermo Fisher | A1517001 |
| DMEM/F-12, GlutaMAX Supplement | Thermo Fisher | 31331093 |
| RPMI 1640 Medium | Thermo Fisher | 21875034 |
| PneumaCult-Ex Plus Medium | Stemcell | 05040 |
| PneumaCult-ALI Medium | Stemcell | 05001 |
| PBS, pH 7.2 | Gibco | 20012019 |
| SuperScript™ First-Strand Synthesis System | Invitrogen | 11904-018 |
| RNeasy Micro Kit | Qiagen | 74004 |
| RNeasy Mini Kit | Qiagen | 74106 |
| LightCycler® 480 SYBR Green I Master | Roche | 04707516001 |
| Alkaline Phosphatase Staining Assay (Red) | ScienCell Research Laboratories | 8288 |
| Alcian blue solution | Merck Millipore | 1016470500 |
| Periodic Acid – Schiff kit | Merck Millipore | 1016460001 |
| 0.4 µm Pore Polyester Membrane Insert (Ø12 mm) | Corning | 3460 |
| TEER device | WPI | EVOM2 Model/ STX2 electrode |
