## Supplemental Table 6 for "Differentiation of human induced pluripotent stem cells into functional airway epithelium"

**Name	Size	Sequences**

GAPDH_F	22	GAC CTG ACC TGC CGT CTA GAA A

GAPDH_R	21	CCT GCT TCA CCA CCT TCT TGA

OCT4_F	22	GGG CTC TCC CAT GCA TTC AAA C

OCT4_R	22	CAC CTT CCC TCC AAC CAG TTG C

NANOG_F	21	TGA TTT GTG GGC CTG AAG AAA

NANOG_R	21	GAG GCA TCT CAG CAG AAG ACA

AFP_F	25	CTA CCT GCC TTT CTG GAA GAA CTT T

AFP_R	22	GAT CGA TGC TGG AGT GGG CTT T

TG_F	20	ACG GTT CCT CGC AGT TCA AT

TG_R	20	GCA GCT TGG AAC ATA GGG GT

CDX2_F	22	ACA GTC GCT ACA TCA CCA TCC G

CDX2_R	22	CCT CTC CTT TGC TCT GCG GTT C

PAX6_F	19	TCT TTG CTT GGG AAA TCC G

PAX6_R	21	CTG CCC GTT CAA CAT CCT TAG

CHGA_F	20	CGG ATC CTT TCC ATT CTG AG

CHGA_R	20	ACC GCT GTG TTT CTT CTG CT

SFTPB_F	22	TCT GAG TGC CAC CTC TGC ATG T

SFTPB_R	22	TGG AGC ATT GCC TGT GGT ATG G

FOXJ1_F	22	GAG ACA GGT TGT GGC GGA TTG A

FOXJ1_R	22	ACT CGT ATG CCA CGC TCA TCT G

MUC5AC_F	20	CAT CTG CCA GCT GAT TCT GA

MUC5AC_R	20	AAG ACG CAG CCC TCA TAG AA

CCSP_F	20	CAT GAA ACT CGC TGT CAC CC

CCSP_R	20	GAT GAC ACG CTG AAA GCT CG
