## Supplemental figure 1 for "Differentiation of human induced pluripotent stem cells into functional airway epithelium"

A

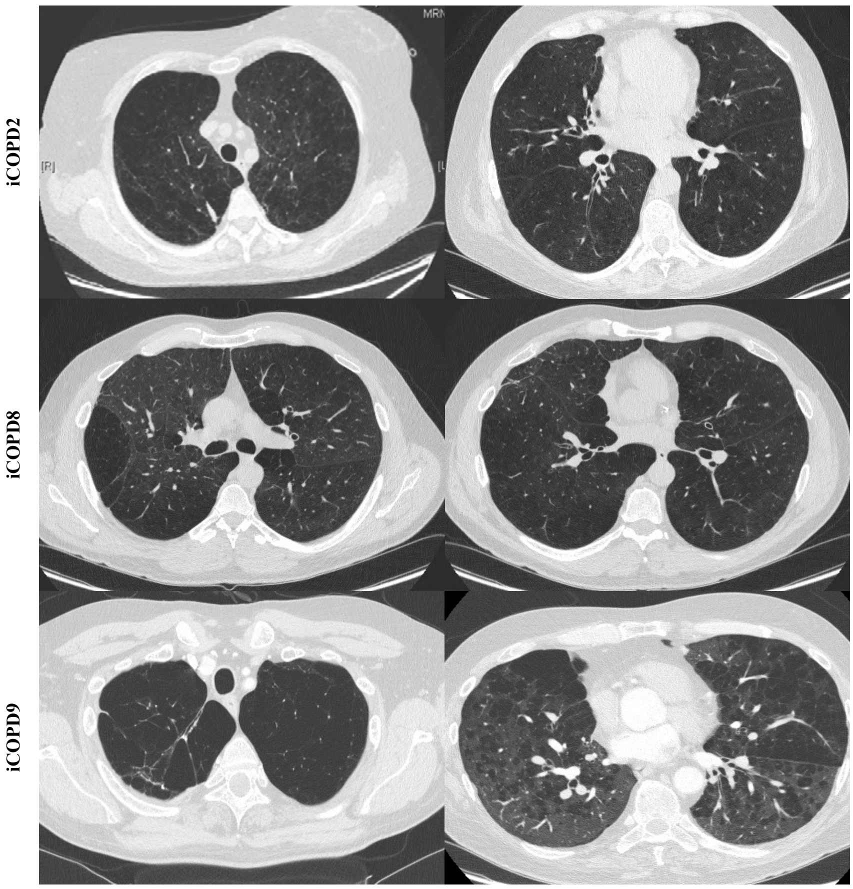


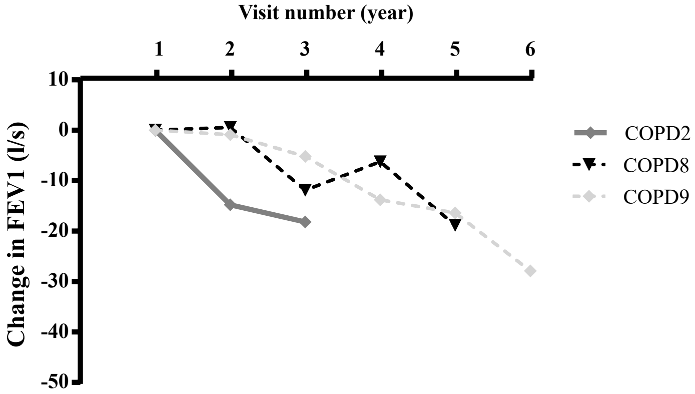


B

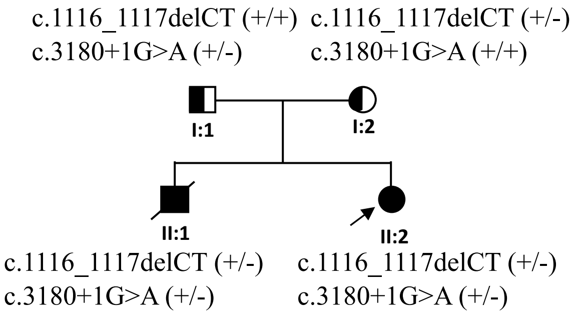


| ID | PCD |
| --- | --- |
| I:1 | No |
| I:2 | No |
| II:1 | TGA, Single Ventricle<br>Pulmonary stenosis<br>Dextrocardia, PCD |
| II:2 | PCD, infertility |

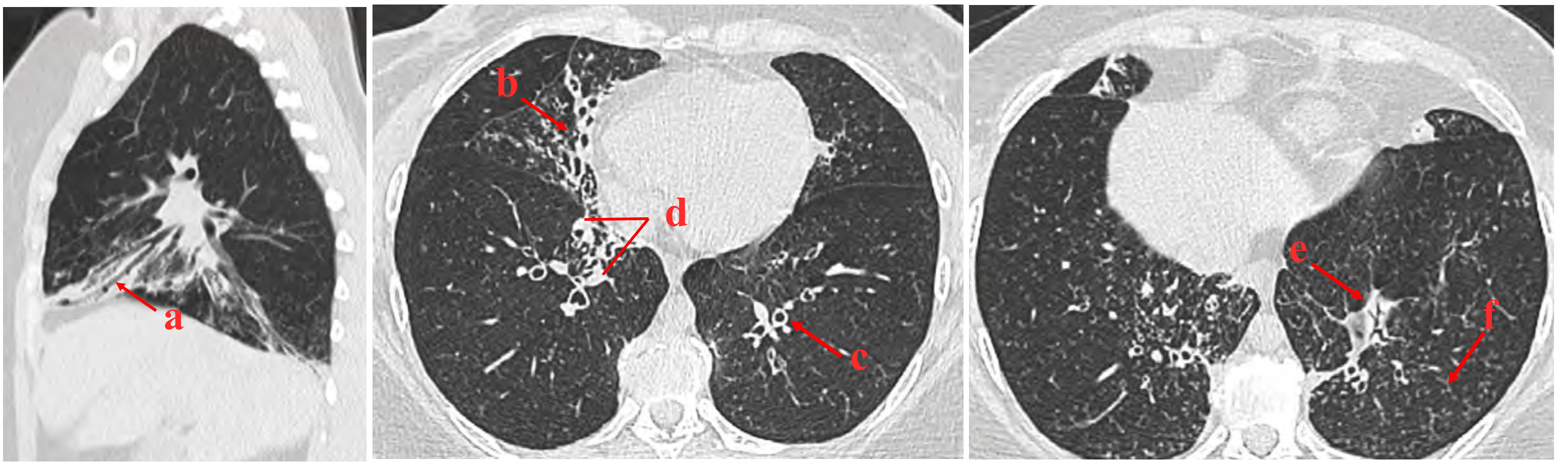
