## Supplementary figures and images for "Differentiation of human induced pluripotent stem cells into functional airway epithelium"

### Supplemental figure 2

A

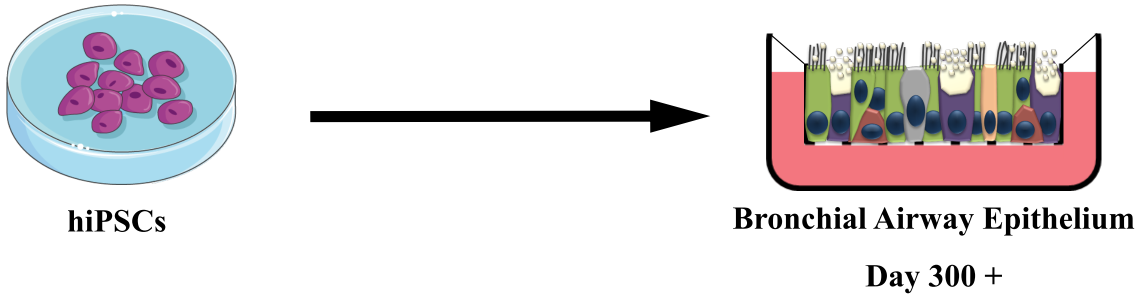


B

iCS- digital test

iCS-digital aneuploidy

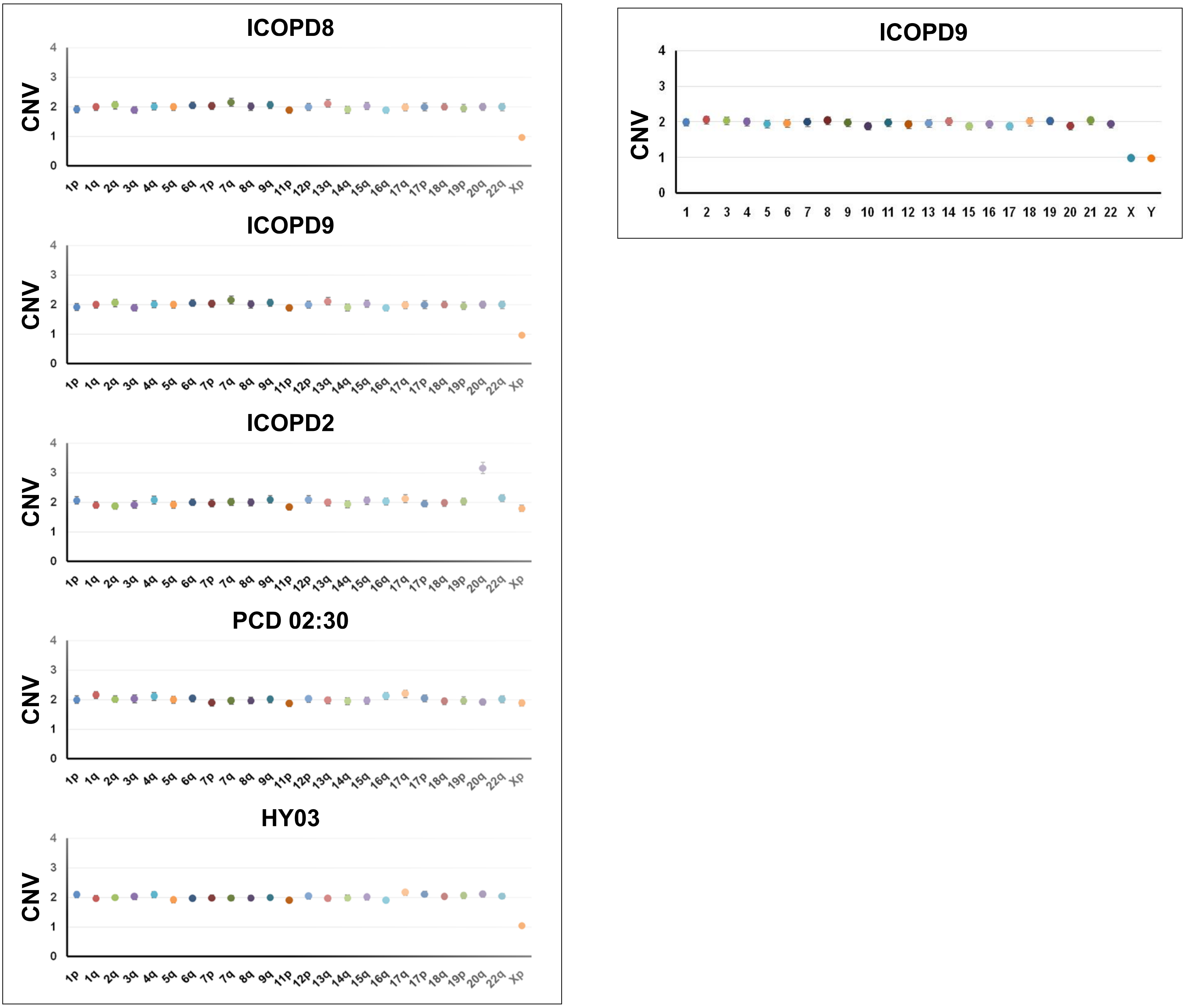

### Supplemental figure 4

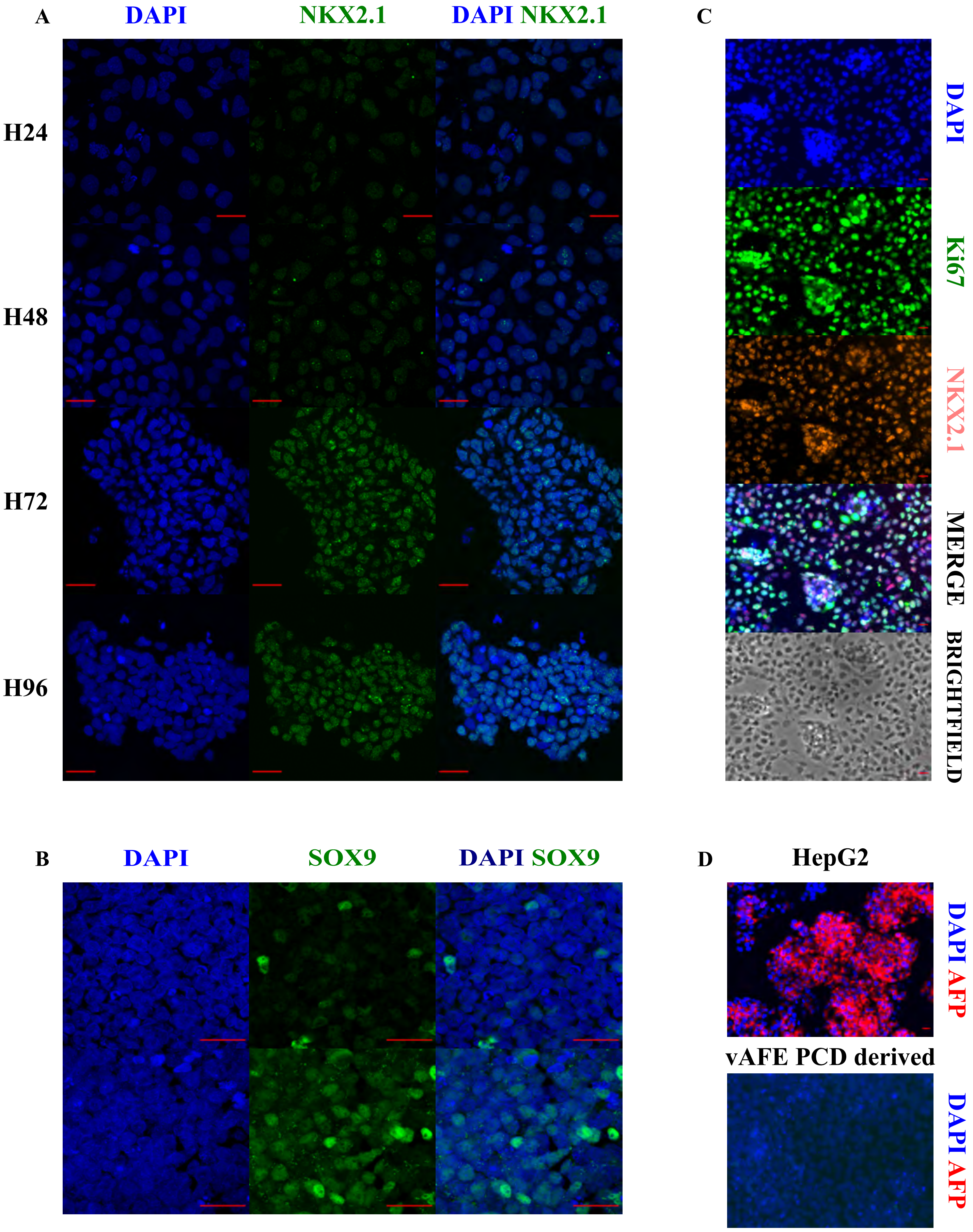

### Supplemental figure 5

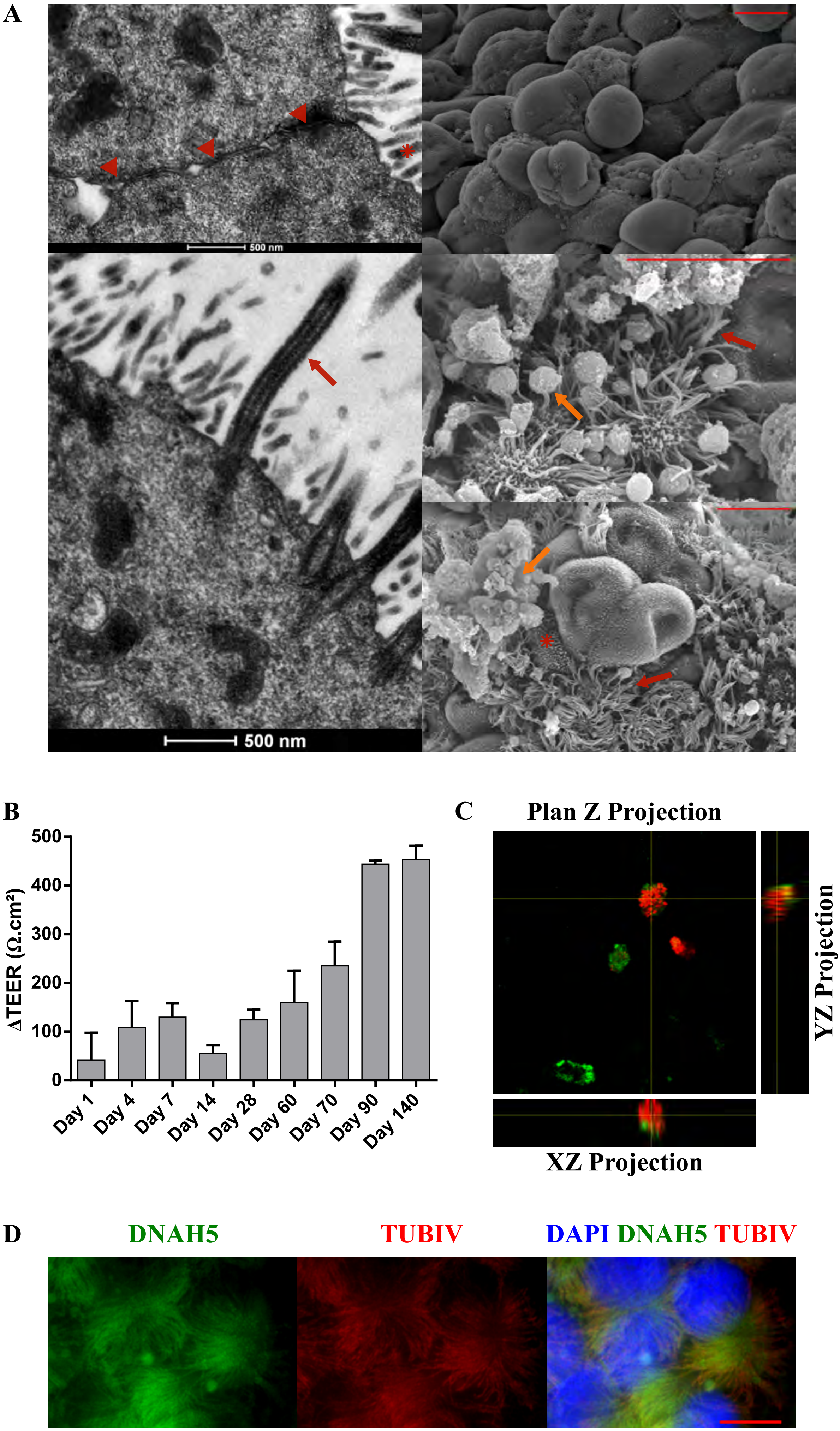
