## Supplemental figure 3 for "Differentiation of human induced pluripotent stem cells into functional airway epithelium"

A - Flow cytometry analysis of CXCR4 at definitive endoderm stage

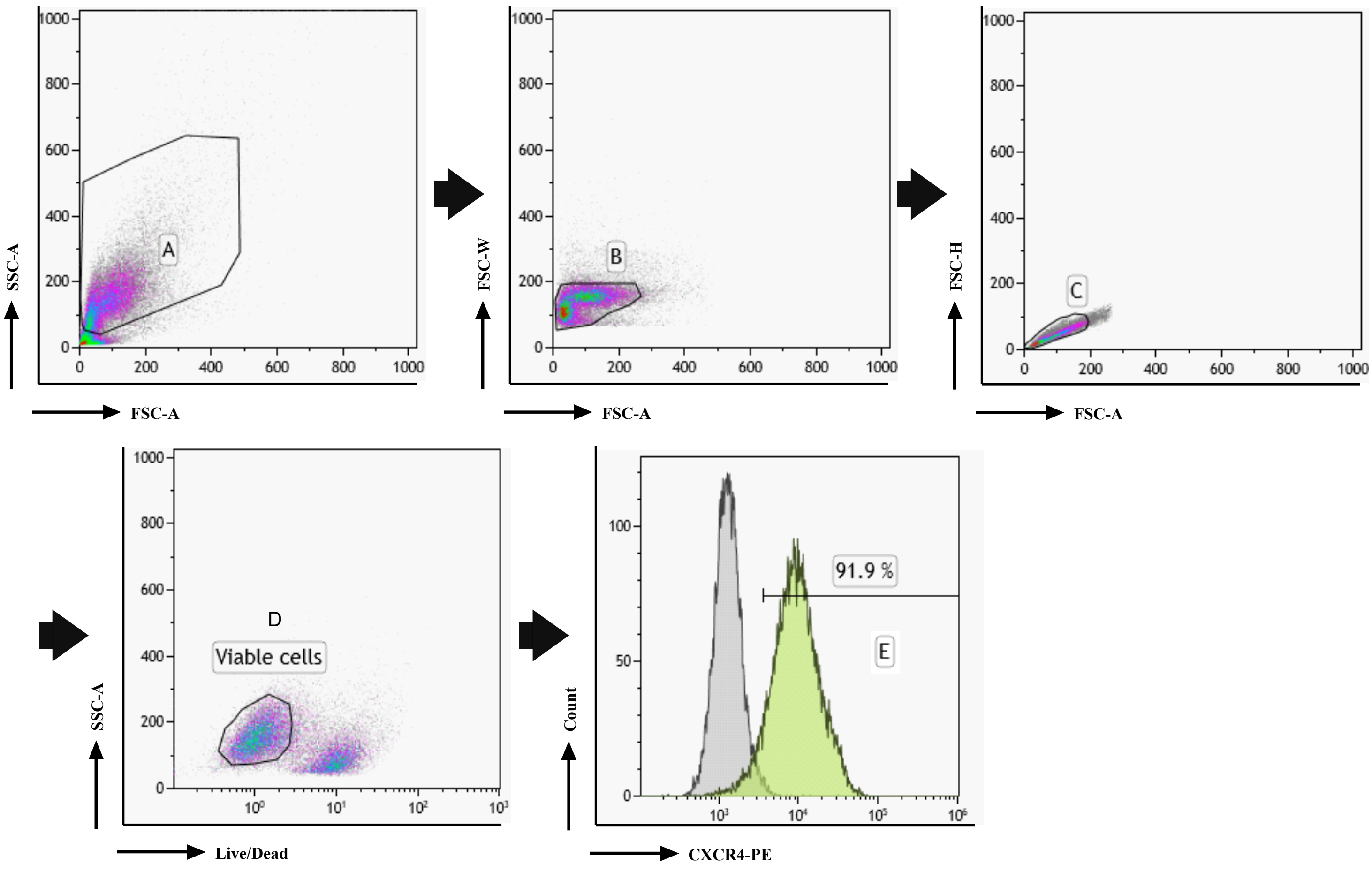


B- Flow cytometry analysis of NKX2.1 at anterior foregut endoderm stage

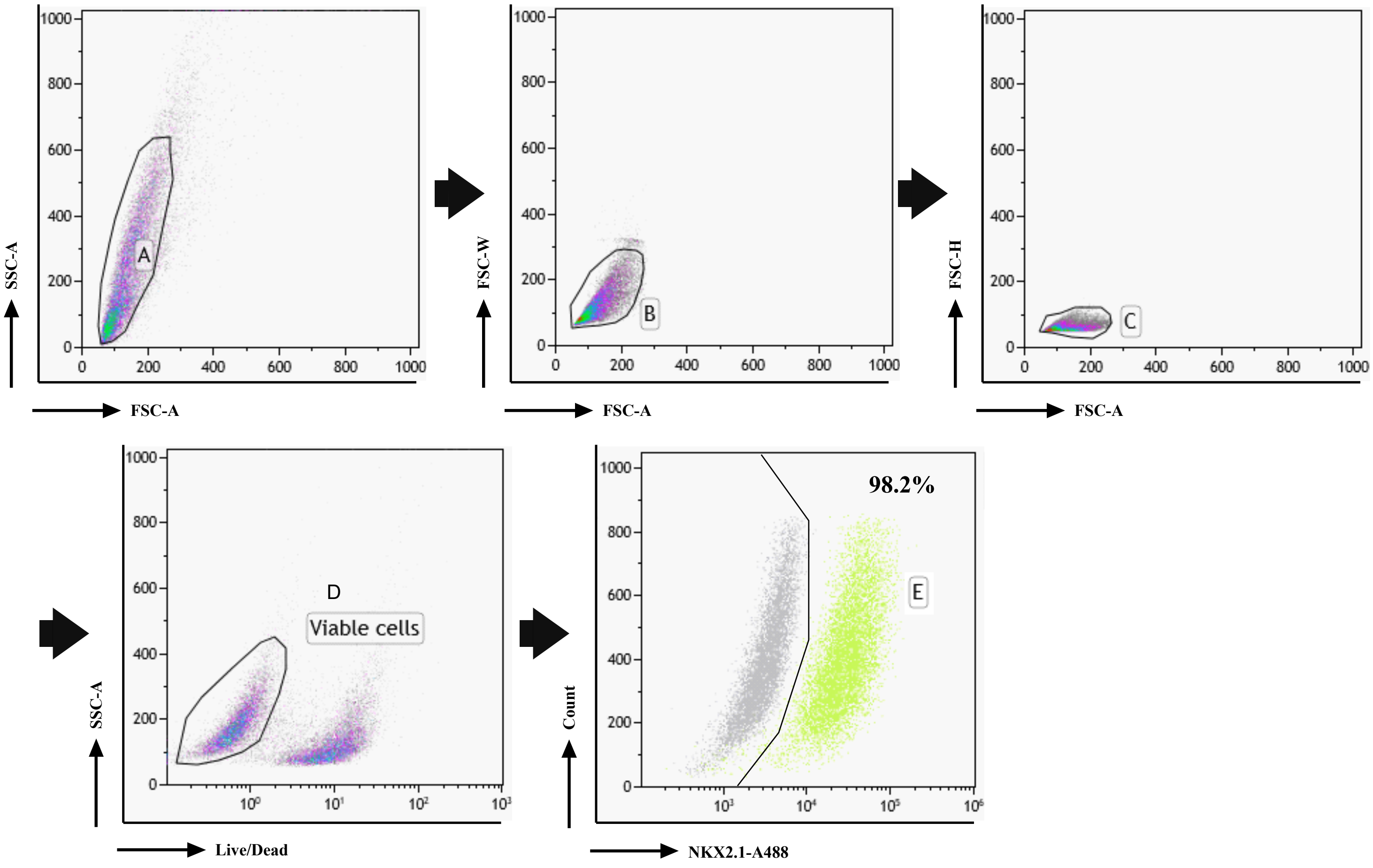
